# Mapping interindividual dynamics of innate immune response at single-cell resolution

**DOI:** 10.1101/2021.09.01.457774

**Authors:** Natsuhiko Kumasaka, Raghd Rostom, Ni Huang, Krzysztof Polanski, Kerstin B. Meyer, Sharad Patel, Rachel Boyd, Celine Gomez, Sam N. Barnett, Nikolaos I Panousis, Jeremy Schwartzentruber, Maya Ghoussaini, Paul A. Lyons, Fernando J. Calero-Nieto, Berthold Göttgens, Josephine L. Barnes, Kaylee B. Worlock, Masahiro Yoshida, Marko Z. Nikolic, Emily Stephenson, Gary Reynolds, Muzlifah Haniffa, John Marioni, Oliver Stegle, Tzachi Hagai, Sarah A. Teichmann

**Author notes:** Corresponding authors, Sarah A. Teichmann Tzachi Hagai.

## Abstract

Common genetic variants modulate the cellular response to viruses and are implicated in a range of immune pathologies, including infectious and autoimmune diseases. The transcriptional antiviral response is known to vary between infected cells from a single individual, yet how genetic variants across individuals modulate the antiviral response (and its cell-to-cell variability) is not well understood. Here, we triggered the antiviral response in human fibroblasts from 68 healthy donors, and profiled tens of thousands of cells using single-cell RNA-seq. We developed GASPACHO (GAuSsian Processes for Association mapping leveraging Cell HeterOgeneity), the first statistical approach designed to identify dynamic eQTLs across a transcriptional trajectory of cell populations, without aggregating single-cell data into pseudo-bulk. This allows us to uncover the underlying architecture and variability of antiviral response across responding cells, and to identify more than two thousands eQTLs modulating the dynamic changes during this response. Many of these eQTLs colocalise with risk loci identified in GWAS of infectious and autoimmune diseases. As a case study, we focus on a COVID-19 susceptibility locus, colocalised with the antiviral OAS1 splicing QTL. We validated it in blood cells from a patient cohort and in the infected nasal cells of a patient with the risk allele, demonstrating the utility of GASPACHO to fine-map and functionally characterise a genetic locus. In summary, our novel analytical approach provides a new framework for delineation of the genetic variants that shape a wide spectrum of transcriptional responses at single-cell resolution.

## Introduction

The innate immune response is a cell-autonomous program that induces an antiviral state in infected and nearby cells, and alerts the immune system of the invading pathogen^1^. Dysregulation of this response can affect a wide range of inflammatory and autoimmune diseases and determine the outcome of infection^2–6^. Common genetic variants have been shown to modulate transcriptional responses to various viral and bacterial stimuli, and to contribute to disease onset and progression^7–11^. Most past gene expression-focused studies of this program are based on bulk RNA-sequencing technologies, which do not fully elucidate the continuous dynamics of transcriptional changes during the innate immune response. Single-cell genomic technologies are powerful approaches to study cell heterogeneity and transcriptional variability across cells^12^. Furthermore, by utilising single-cell RNA-seq (scRNA-seq) profiling of tissues composed of several cell lineages, previous studies have successfully performed genetic association mapping of cell-type specific expression^13–16^.

We here use full-length scRNA-seq of dermal fibroblasts from different human individuals, challenged with immune stimuli. Based on the pseudo-temporal reconstruction of this data, we map the transcriptional variation of the innate immune response at single-cell resolution. This provides the foundation for superimposing human genetic variation onto the transcriptional dynamics of this response. To this end, we develop a novel statistical approach based on a Gaussian process latent variable model^17,18^ called GASPACHO (GAuSsian Processes for Association mapping leveraging Cell HeterOgeneity). This allows us to identify expression quantitative trait loci (eQTL) that manifest at different stages of the response to stimuli.

We find several thousand eQTLs, hundreds of which colocalise with known risk loci of diverse autoimmune and infectious diseases. We perform fine-mapping of the OAS locus, associated with COVID-19, to reveal the imbalanced expression of OAS1 and OAS3 genes during the antiviral innate immune response. We further integrate this data with eQTLs from a COVID-19 patient cohort dataset of Peripheral Blood Mononuclear Cell scRNA-seq, as well as with scRNA-seq data of infected nasal epithelial cells from a COVID-19 patient.

Overall, our study illustrates how coupling single-cell transcriptomics with a cutting-edge statistical approach can identify dynamic effects of human trait-associated genetic variants in different contexts of activation of antiviral innate immunity and, in general, in diverse cellular dynamic processes.

## Results

### Dermal immune stimulation to study antiviral response across cells and individuals

To study the innate immune expression program that is triggered upon viral infection, we exposed primary dermal fibroblasts from 68 donors from the HipSci^19^ to two stimulants: (1) poly(I:C) - a synthetic dsRNA that is rapidly recognized by viral sensors and elicits primary antiviral and inflammatory responses, and (2) Interferon-beta (IFNB), a cytokine that upregulates a secondary wave of response in both infected and bystander cells, and shifts the cells into an antiviral mode, where hundreds of Interferon Stimulated Genes (ISGs) are upregulated in order to contain the infection.

We collected cells exposed to each of the two stimuli after 2 and 6 hours of stimulation (Fig 1a). Following this, single-cell RNA sequencing (scRNA-seq) profiling was performed using a plate-based full-length transcript approach (Online Methods). After QC, 22,188 high-quality cells were obtained across 128 plates, with each plate containing cells from three donors. The donor identity for each cell was inferred from scRNA-seq read data using known genotypes made available by the HipSci consortium (**Extended Data Fig. 1a**; Online Methods). Preliminary analysis showed that our data displays high cell-to-cell variability in gene expression both within and across donors, as observed in previous studies by us and others^20–22^. In fact, our data was confounded by various technical and biological factors, including library preparation in different batches, and cell cycle effects (**Extended Data Fig. 1b**). The complex nature of this data, along with its confounders, motivated us to develop a new approach that reveals the genetic and physiologically-relevant variation, while computationally masking confounding factors.

**Figure. 1.**
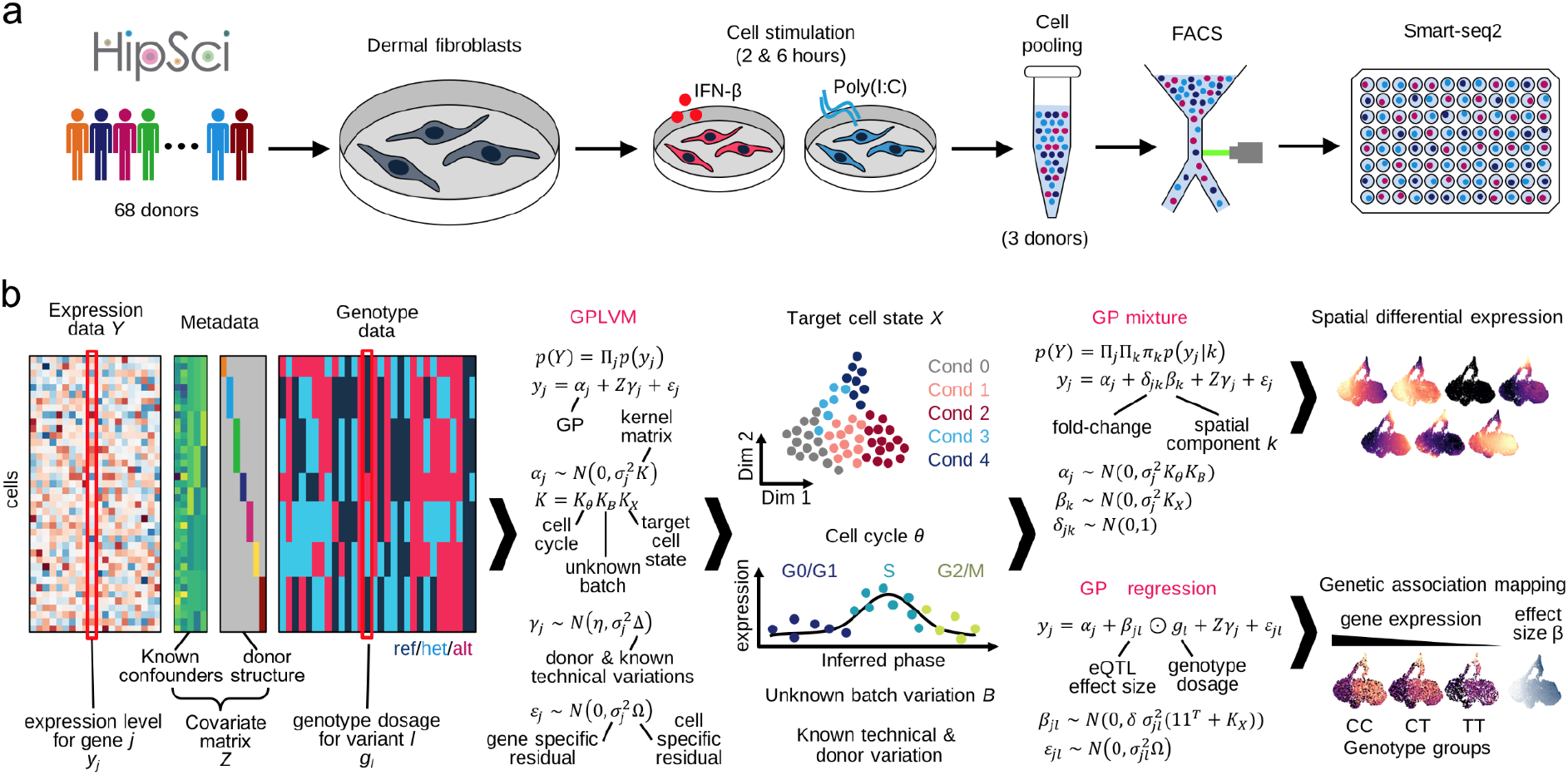
Schematics of experiment and statistical analysis. **a.** Experimental design. **b.** GASPACHO (GAuSsian Processes for Association mapping leveraging Cell HeterOgeneity) framework. Expression data and relevant metadata (know confounding factors) as well as donor (cell line) structure is used to construct a Gaussian Process Latent Variable Model (GPLVM) to extract the target cell state, while dissecting cell cycle effect and other know and unknown technical variability including donor-donor variation. The result of GPLVM is then utilised to the subsequent analyses of spatial differential expression analysis using a GP mixture model and the genetic association mapping using a GP regression model (Online Methods).

### GASPACHO: A novel approach for uncovering cell-state dynamics using a Gaussian Processes

Single-cell transcriptomics (as compared to bulk) enables us to uncover hidden states of complex biological processes, while also requiring regression of technical effects and biological variation that is not of interest (e.g. proliferation). We developed GASPACHO (GAuSsian Processes for Association mapping leveraging Cell HeterOgeneity), which utilises a Gaussian Process latent variable model (GPLVM) to uncover the dynamic cell states of interest, while adjusting periodic cell cycle variation and both known and unknown technical variations simultaneously (**Fig. 1b**; Methods). The use of GPLVM allows us to capture smooth and continuous non-linear trends in gene expression along the latent variables, for which other methods such as the standard linear PCA will not work well.

Although there are other models that utilise GPLVM to study single-cell dynamics^23,24^, the novel aspect of our GPLVM approach is that it explicitly takes into account the donor variation as well as other known confounding effects (such as technical batches) as additional random effect terms (**Fig. 1b**; Methods). These confounders are known to inflate type-I error in downstream analyses, such as in differential expression^25^, leading to false discovery of differentially expressed genes. As detailed below, the model output not only enabled us to look at the architecture of the antiviral response in the cell-state space, but also provided a rigorous statistical framework of (1) spatial differential expression (DE) analysis and (2) genetic association mapping using genotype data obtained from the donor of origin for each cell.

Specifically, the gene expression variation in the target cell-state space was inferred by a Gaussian Process (GP) mixture model in which an additional GP component is introduced into the model to capture hidden spatial DE patterns ^26^ of gene expression in the latent space (**Fig. 1b**; Methods).

The genetic association mapping was also carried out by using a GP regression model in which the effect size of a quantitative trait locus (QTL) was modeled as a GP in the target cell-state space. Here, the additional GP was multiplied by the genotype dosage (the number of alternative alleles for each donor) to capture the gene-environment interaction^27^ (**Fig. 1b**; Methods). Importantly, the eQTL effect is obtained at single-cell resolution, and the model does not require aggregation of single-cell data into pseudo-bulk data, which is a common eQTL mapping strategy. Thus, we can study the effect of genetic variants without losing the continuum of transcriptional dynamics and its spectrum across individual cells. We have implemented the software in R, which is available from github (https://github.com/natsuhiko/GASPACHO).

### Primary and secondary responses of innate immunity

We first applied the GPLVM to adjust for the cell cycle and unknown batch effects in our data (**Extended Data Fig. 1c-d**) and successfully extracted the innate immune state embedded in the data (**Fig. 2a**). We also confirmed that the extracted immune state was orthogonal to cell cycle or the unknown batch variations (**Extended Data Fig. 1e**). We observed two major cell trajectories: one for response to IFN-b from the naive state (x-axis) and the other for response to Poly (I:C) (y-axis).

**Figure 2.**
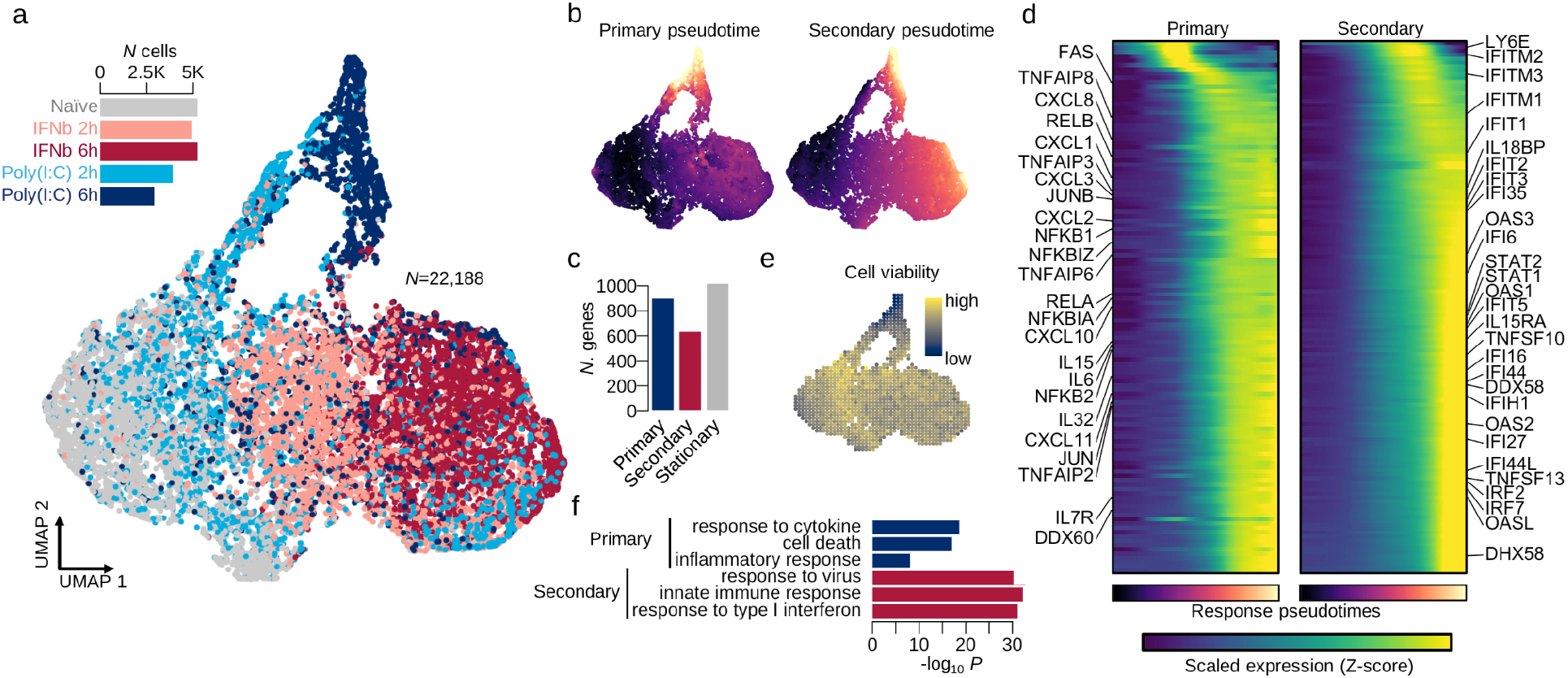
Innate immunity captured by GPLVM and GP mixture model. **a.** UMAP of latent variables capturing innate immune variation between cells. Cells are coloured by the five experimental time points (gray: naive state; pink: IFN-beta 2 hours; brown: IFN-beta 6 hours; blue: poly(I:C) 2 hours; naivy: poly(I:C) 6 hours). **b**. Estimated pseudo time for primary and secondary responses using the GP mixture model. **c**. Barplot shows the numbers of response and stationary genes. **d**. Heatmaps show dynamic gene expression changes along primary or secondary response pseudo time. The pseudo time colour scale corresponds to panel **b**. The expression colour (navy to yellow) shows the magnitude of scaled expression for each gene (Z-score). **e**. UMAP shows a predicted Achilles cell viability using CEVIChE (CEll VIability Calculator from gene Expression) tool (Online Methods). **f**. Barplot shows the enrichment of GO terms for primary and secondary genes.

We then applied the GP mixture model which revealed two independent innate immune responses, the primary response by virus infection and the secondary response for bystander cells due to IFN-b secretion by the infected cells or direct IFN-b stimulation (**Fig. 2b**; **Extended Data Fig. 2a**). Those responses were highly overlapping on the UMAP suggesting those two processes are independently and simultaneously happening in each cell. Interestingly, the primary response was also correlated with the predicted cell viability by CEVIChE (**Fig. 2c**; Online Methods). In total, the GP mixture model discovered 903 and 636 genes upregulated during the primary and secondary responses respectively (hereafter referred to as *primary response genes* and *secondary response genes*), while 1,020 genes were expressed uniformly across all cells in different experimental conditions (referred to as *stationary genes*) (**Fig. 2d**; **Extended Data Fig. 2b**). Many cytokine and chemokine genes were upregulated along the primary response, while interferon stimulated genes (IGSs) were upregulated along the secondary response (**Fig. 2e**). The GO enrichment analysis for the primary and secondary genes clearly demonstrated primary response genes are enriched for cell death and inflammatory response, while secondary response genes are enriched for type I interferon response (**Fig. 2f**; **Extended Data Fig. 2c**).

### Dynamic genetic effect on transcriptional response to innate immune stimuli

We then mapped expression quantitative trait loci (eQTLs) along innate immune response using the GP regression model to assess genetic association in single cell resolution (Online Methods). We discovered 2,662 eQTL genes (local FDR 10%) among 10,748 genes expressed in, at least, 10% of total cells.

In order to demonstrate that we chose a sensible approach, we first compared our GP approach with pseudo bulk approaches and the standard marginal effect approach to confirm that our approach provides both the highest sensitivity and specificity for mapped eQTLs (**Extended Data Fig. 3a-b**; Online Methods).

Among our eQTLs, we found 16% and 13% are primary and secondary response genes (**Fig. 3a, Extended Data Fig. 3c**). Those genes are strongly enriched with the discovered eQTLs (**Fig. 3a, Extended Data Fig. 3d**). We also found the primary response eQTLs are depleted in GTEx fibroblast eQTLs while the stationary eQTLs are enriched (**Fig. 3a, Extended Data Fig. 3e-f**; Online Methods), suggesting our eQTLs are highly context specific.

**Figure 3.**
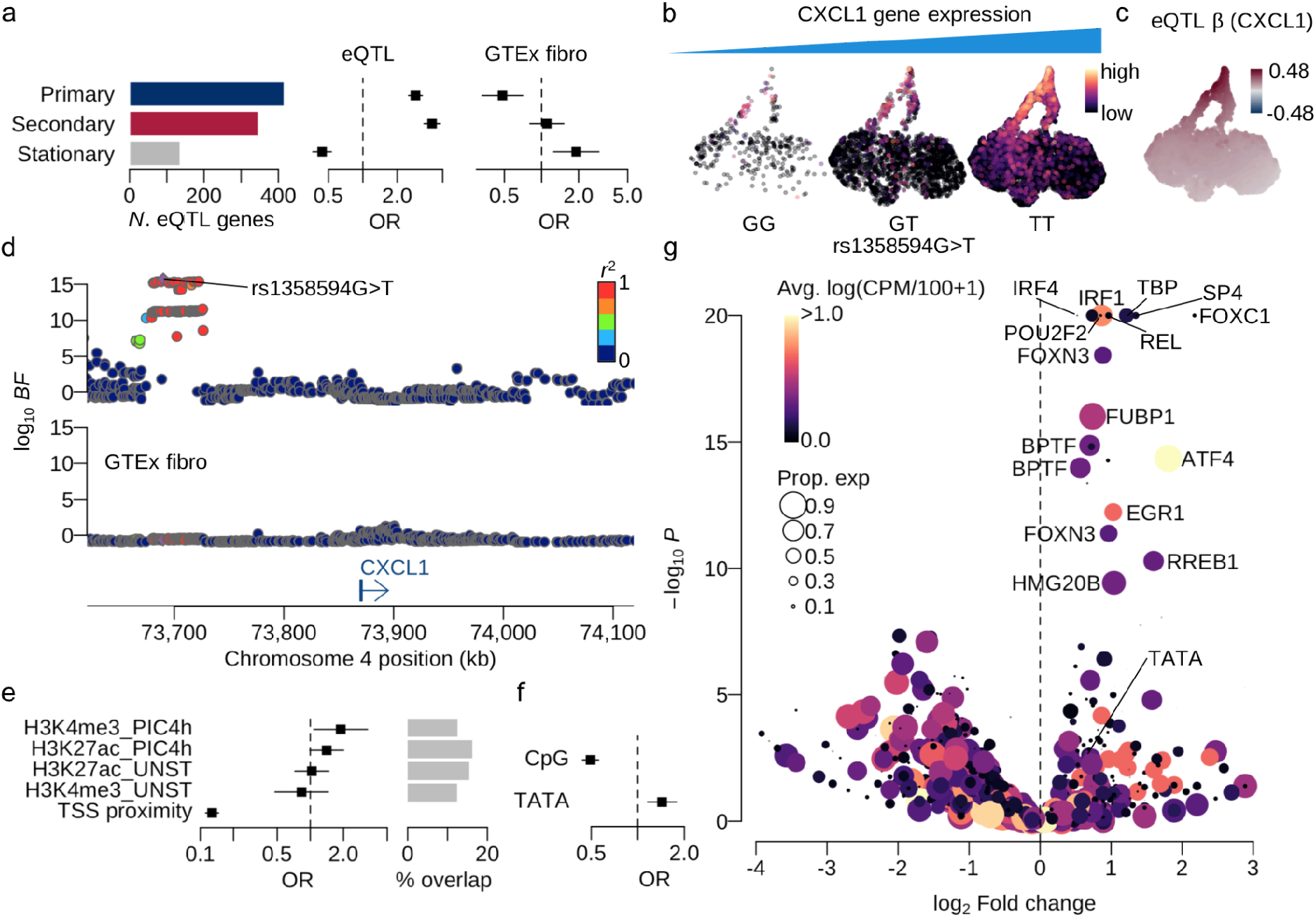
Characteristics of dynamic eQTLs mapped using GP regression. **a**. Barplot shows the numbers of eQTLs (local FDR < 10%) that are primary and secondary response genes or stationary genes. Forest plots show the enrichment of discovered eQTLs for the DE categories and GTEx fibroblast eQTLs. **b**. UMAPs show CXCL1 expression levels stratified by different genotype groups at rs1358594G>T. **c**. UMAP shows the distribution of eQTL effect size (β) at rs1358594. The alternative allele (T) is assessed. **d**. Locus zoom plot of CXCL1 eQTL association Bayes factors around CXCL1 gene (top: in-house data; bottom: GTEx fibroblast eQTL). **e**. Enrichment eQTLs for regulatory regions characterised by ChIP-seq for the two histone modifications (H3K27ac and H3K4me3) under two different conditions (UNST: unstimulated; PIC4h: Poly (I:C) 4 hour stimulation). **f**. eQTL enrichment for genes with TATA box or CpG island between TSS and 100bp upstream. **g**. Enrichment of lead eQTL variants for various transcription factor motifs.

As an example, the CXCL1 gene is a primary response eQTL mostly expressed in later time points of Poly (I:C) stimulated cells (**Fig. 3b**) and the expression level is higher for the alternative allele T at rs1358594 compared with the reference allele G (**Fig. 3c**). This eQTL signal was discovered more than 100Kb downstream of the gene TSS (transcription start site) and only present upon cell stimulation by Poly (I:C), but not in the naive condition as clearly shown in GTEx fibroblast eQTL data (**Fig. 3d**). Note however that this eQTL was discovered in eQTLGen data with tens of thousands of blood samples (**Extended Data Fig. 3g**). This might suggest that the eQTL signal is present in-vivo, although the effect size could be too small to discover with only hundreds of GTEx samples.

We have also performed fine-mapping using epigenetic data (histone modification ChIP-seq of active promoters and enhancers) originating in dsRNA-stimulation of human dermal fibroblasts^22^ (Methods). We identified more than 10% of putative causal eQTL variants which are found in each ChIP-seq peak and enriched for promoter peaks of Poly (I:C) stimulated cells characterised by H3K4me3 antibody (**Fig. 3e**). Note that our eQTLs were also strongly enriched around TSS, thereby the number of eQTLs was reduced by 34% every 100Kb further away from TSS (**Fig. 3e**).

We next tested whether promoter architecture affects the variability between individuals. It was previously shown by us and others^22,28^ that genes containing TATA-boxes in their promoters tend to vary more in transcription between species, conditions and between individual cells responding to immune stimulus, whereas promoters containing CpG-Islands (CGIs) tend to vary less and be transcriptionally more homogenous. We observe that genes with TATA-containing promoters are 1.4 times more highly enriched with eQTLs in comparison with genes with CGI-containing promoters (**Fig. 3f**).

Using the fine-mapped eQTL variants using ChIP-seq annotations, we finally examined which transcription factor motifs were disrupted by the lead eQTL variants (Methods). We found interferon regulatory factor 1 and 4 (IRF1 and IRF4) as well as REL and ATF4 were significantly enriched (**Fig. 3g**). An example of putative TF binding disruption was discovered in RTP4 eQTL, where the alternative allele of a promoter flanking eQTL variant (rs62292793T>A) may disrupt an IRF1 motif that significantly reduces putative TF binding affinity, which subsequently downregulates the RTP4 expression (**Extended Data Fig. 3e**). Furthermore, the TATA-motif is also found to be disrupted by eQTL variants, further suggesting the importance of TATA-regulation in modulating the response and its variability among individuals. *[mention TATA and Landry’s Science 2008 in Discussion]*

### Innate immune response eQTLs are colocalised with autoimmune and infectious diseases

One of the advantages of eQTL mapping is to uncover the target genes and related cell states at each genetic locus implicated by genome-wide association studies (GWAS) of common complex traits. We here tested for colocalization of our eQTLs with risk loci from 701 GWAS with 5 or more genome-wide significant loci, of which 112 were broadly immune related, including autoimmune and chronic inflammatory diseases such as Crohn’s disease and infectious diseases such as COVID-19 (Online Methods). We discovered 3,132 unique gene-trait combinations with the posterior probability of a single shared causal variant between an eQTL and a GWAS locus greater than 0.5. The combinations consisted of 495 different GWAS traits and 823 unique genes. We observed an excess of colocalised eQTLs for immune related traits over non-immune traits (**Fig. 4a**; *P*=2.0×10^−5^; Online Methods), likely reflecting the known involvement of innate immunity in each of the disease pathologies.

**Figure 4.**
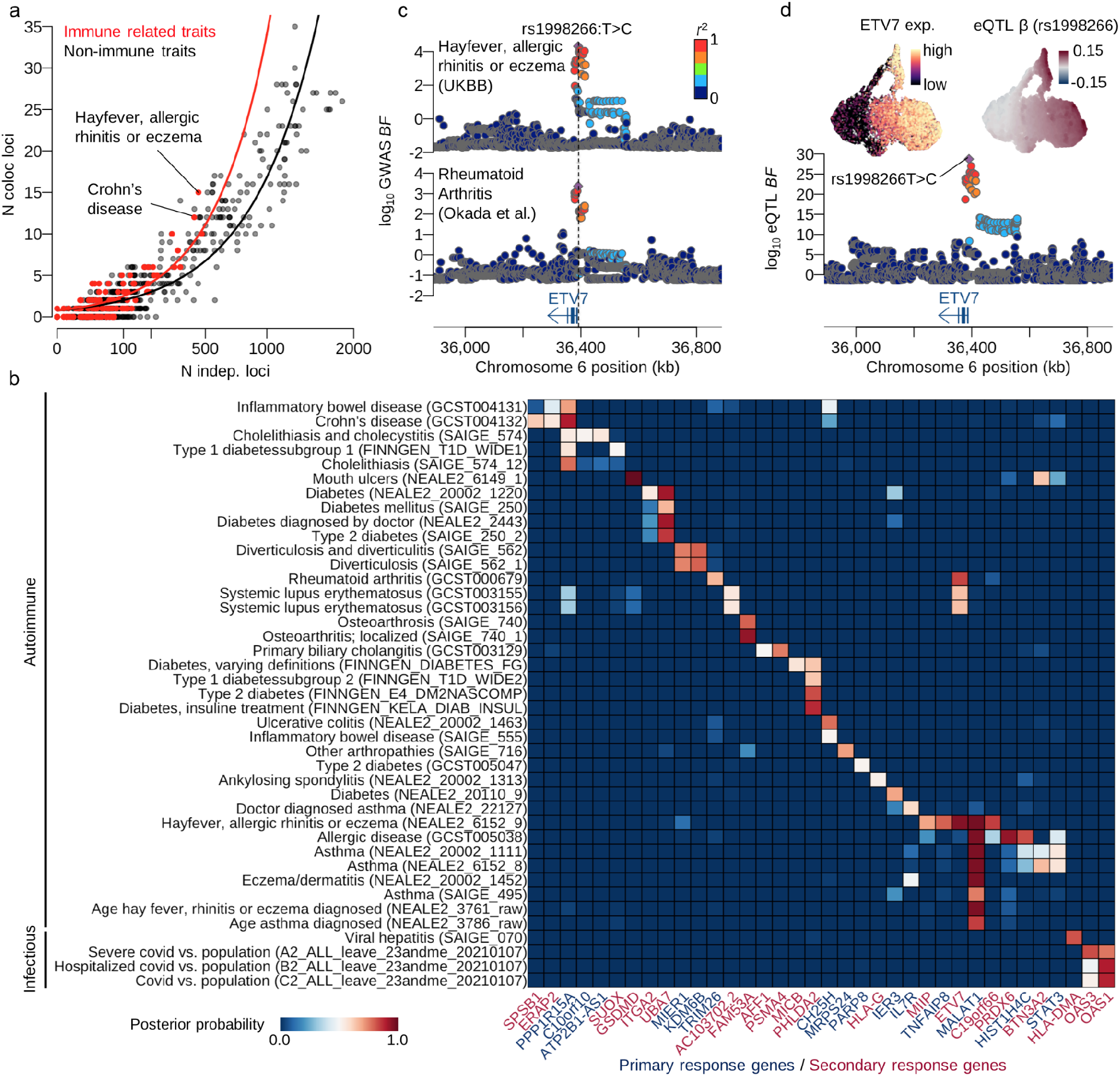
GWAS colocalisation. **a**. Scatter plot shows the number of colocalised eQTLs with posterior probability greater than 0.5 (y-axis) against the number of independent loci where GWAS and eQTL do not share the putative causal variant (Online Method). **b**. Heatmap shows the posterior probability of colocalisation between eQTLs and GWAS loci. Only primary (coloured by navy) and secondary (coloured by red) response genes were shown. **c**. Locus zoom plots show the association of rheumatoid arthritis and hayfever, allergic rhinitis or eczema around ETV7 gene. Points are coloured by the LD index (*r*^2^) with the GWAS index variant rs1008266T>C. **d**. UMAPs show the scaled ETV7 expression and the eQTL effect size (β) at the lead eQTL variant rs1998266T>C. Locus zoom plot shows the eQTL association for ETV7.

We discovered 36 primary and secondary response eQTLs that were specifically colocalised with 41 autoimmune and infectious diseases, some of which were colocalised with multiple traits (**Fig. 4b**). For example, we detected an eQTL for the ETV7 gene which produces a transcription factor in the ETS family and plays a key role in hematopoiesis ^29^. The eQTL was colocalised with rheumatoid arthritis and hayfever, allergic rhinitis or eczema (**Fig. 4c**). The gene is an ISG and the expression is upregulated during secondary response (**Fig. 4d**). The lead eQTL variant (rs1998266T>C) is shared with the GWAS traits, whose alternative allele C upregulates gene expression in stimulated conditions and also increases the risks of those GWAS traits (**Fig. 4d**). The alternative allele C also modifies the binding motif of the transcription factor ATF6 putatively bound at the promoter region of ETV7, thereby potentially increasing the expression level (**Extended Data Fig. 4**).

### Fine-mapping OAS1 eQTL associated with SARS-CoV-2 infection

In conjunction with the fibroblast system, we used two additional *in vivo* systems (**Fig. 5a**) to further finemap the 12q24.13 (OAS) locus which was reported in a genome-wide association study of reported SARS-CoV-2 positive-infected individuals against population controls^30^ (index SNP: rs10774671G>A). The locus is colocalised with the OAS1 eQTL in fibroblasts with a posterior probability of 0.89 (**Fig. 4b**, **Fig. 5b**). OAS1 is a secondary response gene, and is highly expressed upon IFN-b (at 2h and later) and Poly (I:C) stimulation (at 6h) (**Fig. 5c**). The alternative allele A of rs10774671 down-regulates the expression level (**Fig. 5d**).

**Figure 5.**
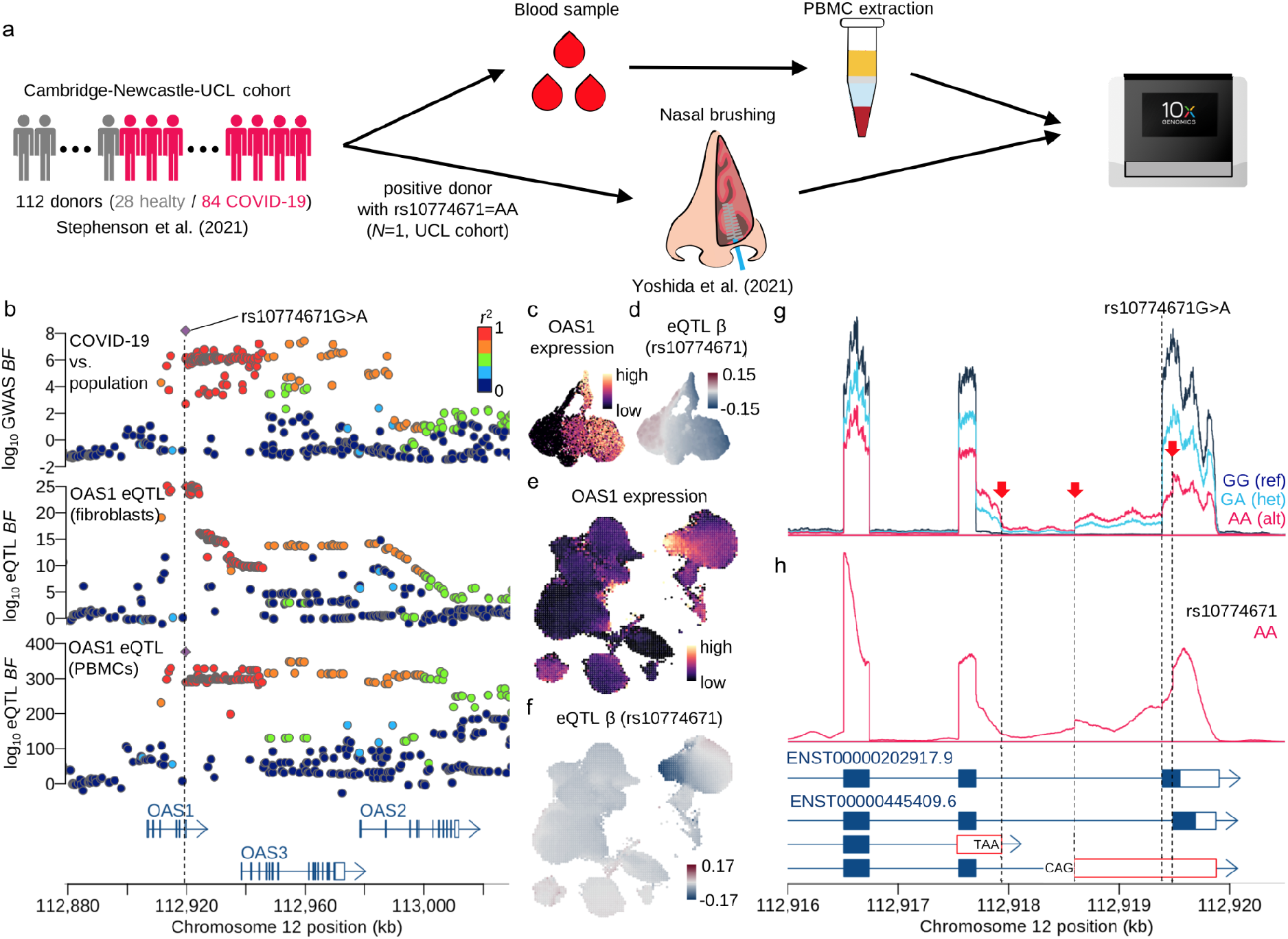
Fine-mapping OAS locus using three different systems. **a**. Schematic of *in vivo* system, COVID-19 study of PBMCs and a nasal brushing to confirm the splicing QTL association in OAS1 gene. **b**. Locus zoom plots show the COVID-19 GWAS (COVID-19 vs population) association Bayes factors as well as eQTL associations of OAS1 gene in fibroblasts and PBMCs. **c**. UMAP shows the expression levels of OAS1 gene in fibroblast. **d**. UMAP shows the eQTL effect size of OAS1 gene at rs10774671G>A. **e.** UMAP shows OAS1 expression level in PBMCs. **f**. UMAP shows the eQTL effect size of OAS1 gene at rs10774671G>A in PBMCs. **g.** Sequencing coverage depth around the splicing variant rs10774671G>A which creates three different isoforms, two of which are not annotated in Ensembl 90. Single cell RNA-seq reads in fibroblasts were aggregated and stratified by the three different genotype groups (GG: reference homozygote; GA: heterozygote; AA: alternative homozygote). **h.** The sequencing coverage depth of epithelial cells for one COVID-19 positive patient with alternative homozygote AA at rs10774671 (Online Methods).

We investigated our recently published PBMC scRNA-seq data^31^ obtained from 112 donors, including 84 COVID-19 positive individuals, as an independent in vivo validation of OAS1 colocalisation with COVID-19 GWAS (Online Methods). There are 18 major blood cell types annotated in this data set (**Extended Data Fig. 5a**), of which myeloid cells and certain T cell subtypes show higher expression of the secondary response genes where the secondary innate immune response is inferred from our fibroblast data (**Extended Data Fig. 5b**; Online Methods). As expected, OAS1 is highly expressed as a secondary response gene in PBMCs (**Fig. 5e**) and we confirmed that OAS1 is also a strong eQTL in PBMCs and colocalises well with the COVID-19 GWAS locus with the posterior probability of 0.99 (**Fig. 5b**). The GWAS index variant rs10774671G>A is the lead eQTL variant in PBMCs whose alternative allele A is strongly negatively correlated with OAS1 expression. This is especially clear in CD16+ monocytes, among other immune cell types (**Fig. 5f**, **Extended Data Fig 5a**).

The index SNP rs10774671 is known to be a splicing QTL^32^ that disrupts the splicing motif right next to the last exon of OAS1 gene (**Fig. 5g**). In our fibroblast data, this variant also increased the intron expression between the last two exons and created the three different isoforms, all of which are known to cause impaired OAS1 protein expression^32^. These alternative isoforms are also observed in nasal epithelial cells in the nasal brushing sample of a COVID-19 positive patient with the alternative homozygous genotype (**Fig. 5a, h**). The cells from this patient contain SARS-CoV2 viral reads, although the same cells express OAS1 and SARS-CoV2 at relatively low levels^33^. Since the alternative allele A of rs10774671 is also the risk allele in COVID-19 GWAS, this implies that impaired OAS1 RNA, and hence, protein expression may cause dysregulation of SARS-CoV-2 virus RNA degradation and clearance in host cells.

## Discussion

In this work we developed GASPACHO - a novel statistical framework that allows for the first time, to infer how genetic variants affect the trajectory of gene expression over a dynamic process such as a stimulation time course across individual cells.

Using GASPACHO, we integrated scRNA-seq data from fibroblasts from 68 donors stimulated by innate immune stimuli, and obtained a low dimensional gene expression space representing the response dynamics across stimulated cells. This procedure also provides us with a map of interindividual transcriptional variation at single-cell resolution, which were also enriched for regulatory regions (such as TF binding sites) profiled during fibroblast stimulation^22^. This approach discovered 2,662 eQTL loci, of which 823 were colocalised with one or more GWAS associated loci of autoimmune and infectious diseases including COVID-19 at the OAS locus.

In conjunction with the OAS1 eQTL, OAS3 eQTL in fibroblasts was also colocalised with COVID-19 GWAS (PP=0.53) (**Fig. 4b**, **Extended Data Fig. 6a**). Because OAS1 and OAS3 are both interferon stimulated genes, the expression patterns of OAS1 and OAS3 along the innate immune response trajectory are very similar (**Fig. 5c**, **Extended Data Fig. 6b**). However the eQTL effect direction was opposite for the two genes (**Fig. 5d**, **Extended Data Fig. 6c**): OAS1 gene expression is downregulated by the alternative allele of the index SNP rs10774671G>A, whereas the expression level of OAS3 is upregulated by the alternative allele. According to the risk allele of COVID-19 GWAS, OAS1 is more likely to be the putative causal gene for SARS-CoV-2 infection because the COVID-19 risk allele matches the impaired protein expression of OAS1 (**Extended Data Fig. 6d**).

Fibroblasts are not the primary cellular target of SARS-CoV-2 infection in most tissues (excepting certain decidua fibroblasts), as they do not express ACE2 (only 8 cells among 22,188 cells express ACE2 in our dermal fibroblast dataset). However, our findings suggest that OAS1 expression can be modulated by a common splicing variant, which occurs in viral target cell types, such as nasal epithelial cells. In these cells, the splicing variant will likely directly influence the efficacy of viral RNA clearance and degradation in the host.

The GPLVM implemented in GASPACHO was applicable for more than 20K cells, the current implementation in R is not scalable for hundreds of thousands of cells. In order to overcome this issue, we need a cutting-edge Bayesian inference technique, such as the stochastic variational inference implemented on modern GPU machines.

In summary, our study demonstrates how an *in vitro* system combined with single-cell RNA transcriptomics, allows us to chart the transcriptional landscape of complex innate immune responses. Our single-cell datasets combined with the Gaussian process based approach shed light on the common genetic basis of autoimmune and infectious diseases during this challenging period of the COVID-19 pandemic.

## Acknowledgements

N.K., R.R., T.H. and S.A.T. were supported by Wellcome Sanger core funding (WT206194) and the Human Induced Pluripotent Stem Cell Initiative. R.R was supported by the BBSRC Doctoral Training Programme. F.J.C.-N. and B.G. were supported by Wellcome (206328/Z/17/Z) and MRC (MR/S036113/1). P.A.L was supported by the Evelyn Trust (20/75) and the UKRI/NIHR through the UK Coronavirus Immunology Consortium. K.B.W. acknowledges funding from University College London, Birkbeck MRC Doctoral Training Programme. M.Z.N. acknowledges funding from the Rutherford Fund Fellowship allocated by the MRC and from the UK Regenerative Medicine Platform 2 (MR/5005579/1). M.Z.N and K.B.M have been funded by the Rosetrees Trust (M944) and from Action Medical Research (GN2911). We would like to thank Michael D. Morgan for careful reading of the manuscript and data sharing.

## Author contribution

R.R., O.S., T.H. and S.A.T. designed the experiments of fibroblast stimulation and single-cell sequencing. R.R., S.P., R.B., C.G., S.N.B. and T.H. performed the experiments. N.K. and S.A.T. developed the analysis method. N.K., R.R., N.H. and T.H. analysed the data. N.I.P. provided the HipSci genotype data. J.S. and M.G. provided the Open Targets GWAS summary statistics. K.B.M., P.A.L., F.J.C.-N., B.G., J.L.B., K.B.W., M.Y., M.Z.N., E.S. and M.H. performed genotyping of the PBMC samples. G.R., M.H., and J.M. provided the processed single cell RNA-seq data of the PBMC samples. M.Y., K.B.W., M.Z.N. and K.B.M. provided the data from the COVID-19 nasal brushing sample. N.K., K.P. and N.H. analysed the COVID-19 nasal brushing data.

## Methods

### Cell culture and stimulation

Primary dermal fibroblast cells from the Human Induced Pluripotent Stem Cell Initiative (HipSci; http://www.hipsci.org/) were used. The cells were derived from healthy individuals spanning a range of ages and both genders. Following a similar protocol used in our previous work^22^, cells were cultured in DMEM (high glucose, pyruvate - Life Technologies), with 10% FBS, GlutaMAX and 1% penicillin-streptomycin. In each experimental batch, we cultured in parallel cells from three different individuals. Cells were split the day before the experiment into separate wells and on the day of experiment were stimulated with either dsRNA (0.5 *μ*g/ml rhodamine-conjugated poly(I:C), transfected with 1 *μ*l/mL lipofectamine 2000, for 2 or 6 hours), 1000 U/ml human recombinant IFN-B, for 2 or 6 hours, or left untreated.

After the relevant period of time, cells were trypsinised and resuspended in PBS. Samples from the three individuals with the same treatment were mixed (for example, ‘unstimulated’ cells from the three donors would be pooled together). The primary aim of this mixing step was to reduce downstream experimental variability between donors, while simultaneously streamlining the collection stage.

### Sorting and single-cell library preparation

Cells were sorted on a Becton Dickinson Influx into 96-well plates containing 2 *μ*l/well lysis buffer. Single cells were sorted individually (using FSC-W vs FSC-H), and apoptotic cells were excluded using DAPI. Rhodamine positive cells were selected in the poly(I:C) treatments. Reverse transcription and cDNA amplification were performed according to the SmartSeq2 protocol (Picelli et al., 2014), and library preparation was performed using an Illumina Nextera kit. Samples were sequenced using paired-end 75bp reads on an Illumina HiSeq 2500 machine.

### Cell viability prediction

The cell viability was predicted by the web based tool called CEVIChE (CEll VIability Calculator from gene Expression; https://saezlab.shinyapps.io/ceviche/). Because the tool is designed for bulk RNA-seq data, we aggregated gene expression levels for neighbouring cells based on the UMAP in Fig. 2a. We constructed 30 × 30 equispaced grids and took geometric means of log CPM values within each grid.

### Smart-seq2 data preprocessing and quality control

We preprocessed the Smart-seq2 data exactly the same as in Kumasaka et al.(REF). We used demuxlet (https://github.com/statgen/demuxlet) to identify the genetic origin of each cell as well as to remove doublets using the genotype data from HIPSCI (see below).

### Genotype data

We obtained the SNP genotype data from HipSci^19^. We converted the genome coordinate from hg19 to GRCh38 using CrossMap (version 0.5.2; http://crossmap.sourceforge.net/).

### Gaussian Process Latent Variable Model

The GASPACHO framework incorporated a GPLVM as a core model to estimate the latent variables and model parameters subsequently used in the differential expression analysis and eQTL mapping. We assumed the gene expression vector *y*_*j*_ = (*y*_*ij*_; *i* = 1, …, *N*)^*T*^ for the gene *j* across *N* cells is independently drawn from

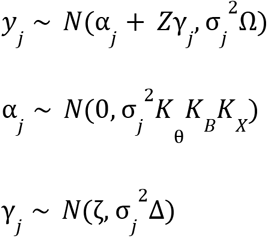

where α_*j*_ is a baseline GP governed by three different kernel matrices, periodic kernel matrix*K*_θ_ for the cell cycle state (θ) and two other squared exponential kernel matrices *K*_*B*_ and *K*_*X*_for unknown batch effects (*B*) and the target cell state (*X*), respectively. Here *Z* is a design matrix for the known covariates, such as donor and sequencing plates (**Fig. 1b**), and γ_*j*_ is a random effect to adjust the known confounding effects whose mean and variance were defined by ζ and the diagonal matrix Δ shared across all genes *j* = 1, …, *J*. The residual expression was determined by the gene specific residual variance *σ*_*j*_^2^ and the cell specific residual variance Ω = *diag*(ω_*i*_; *i* = 1, …, *N*). Note that the variances of the GP and the random effect were properly scaled by multiplying the gene specific residual variance *σ*_*j*_^2^.

The model parameters {Δ, Ω, Σ, ζ} and the latent variables {θ, *B*, *X*} were inferred by maximising the marginal likelihood

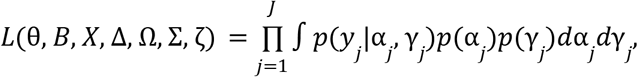

where Σ = *diag*(σ_*j*_^2^; *j* = 1, …, *J*). We used the L-BFGS algorithm with the analytic gradient of the likelihood function with respect to the parameters and the latent variables. In reality, the kernel matrices are not tractable for large *N*, we computed the Titsias bound using the sparse GP ^18^ to approximate the above likelihood. See Supplementary Note for more details.

### Gaussian Process mixture model for spatial differential expression analysis

We employed a GP mixture model to perform differential expression analysis in the target cell state space defined by *X* which was estimated by the GPLVM. Specifically, we introduced one extra GP β_*k*_ for the *k*th differential expression group (*k* = 1, …, *K*) to which a gene *j* belongs:

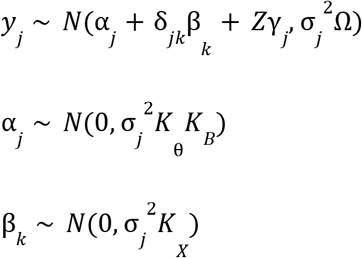

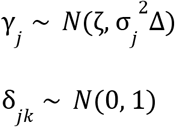

Here the effect size of the GP was properly scaled by a coefficient δ_*jk*_ to allow the GP to be both positively or negatively correlated with the gene expression. The model parameters {Δ, Ω, Σ, ζ} and the latent variables {θ, *B*, *X*} were replaced by the estimated values by the GPLVM. Then we maximised the likelihood of a finite mixture of GPs:

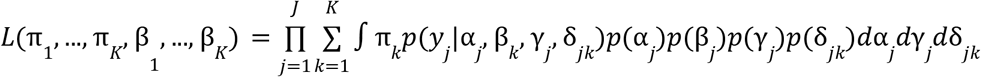

with respect to π_*k*_ and β_*k*_ for *k* = 1, …, *K*. Note that the number of total mixture components *K* is fixed in the current implementation and *K*=3 in the fibroblast data. Again we used the sparse approximation to make the likelihood tractable (see Supplementary Note for more details).

For the pseudotime analysis, we computed the posterior mean **E**[β_*k*_] of the GP for the *k*th component, which provided the underlying cell state regarding the primary and secondary innate immune responses.

### Gaussian Process regression for association mapping

We employed a GP regression model to map eQTLs in the target cell state space defined by *X* which was estimated by the GPLVM. Specifically, we introduced one extra GP β_*jl*_ for the gene *j* multiplied by the *l*th genetic variant *g*_*l*_ = (*g*_*l1*_, …, *g*_*lN*_)^*T*^ whose *i*th element *g*_*li*_ are alternative allele dosages for the individual *i* as a gene environment interaction:

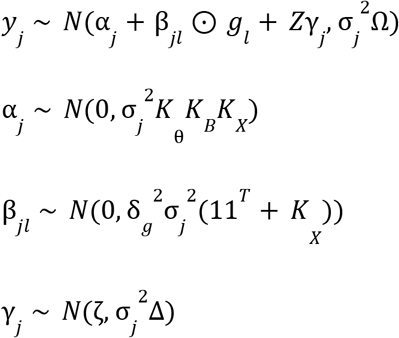

Here the eQTL effect size was properly scaled by a coefficient δ_*g*_ to allow for controlling the genetic contribution on the expression level. Here the model parameters {Δ, Ω, Σ, ζ} and the latent variables {θ, *B*, *X*} were replaced by the estimated values by the GPLVM. The Bayes factor of genetic association can be obtained by:

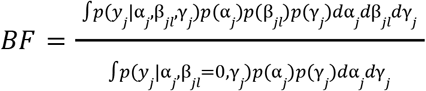

where we set δ_*g*_ = 0. 1 (see Supplementary Note for more details).

### Hierarchical model for eQTL discovery and enrichment analysis using genomic annotations

We tested genetic variants whose minor allele frequency is greater than 0.05 in 1Mb cis regulatory window centred at each gene TSS. In order to control the false discovery rate in a Bayesian framework, we used the hierarchical model^34^ to obtain the posterior probability that a gene is an eQTL as well as the posterior probability that a variant is an eQTL variant within the cis window.

The model allows incorporating various genomic annotations in the gene-level and variant-level as demonstrated previously^34^. We used the ChIP-seq peak annotations obtained from Hagai et al.^22^ in conjunction with TSS proximity to estimate the contribution of epigenetic information to the eQTL variant discovery.

### eQTL enrichment in differentially expressed genes

Any enrichment analysis was carried out based on posterior probability *Z*_*j*_ that the gene *j* is an eQTL obtained from the hierarchical model. We then computed a 2×2 table using a corresponding binary annotation *X*_*j*_ (e.g., if the gene *j* belongs to some annotation, e.g., TATA gene, then *X*_*j*_ = 1 otherwise *X*_*j*_ = 0) or alternatively the posterior probability *X*_*j*_ ∈ [0, 1] that the gene *j* is a DE gene (one of multiple DE gene categories defined above), such that

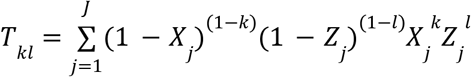

for *k*, *l* = 0, 1. From the table *T*, we computed the log odds ratio 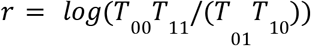 and its standard error *Var*(*r*) = (1/*T*_00_ + 1/*T*_01_ + 1/*T*_10_ + 1/*T*_11_) to perform hypothesis testing. The confidence interval of the log odds ratio was given by 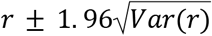.

### eQTL sharing with GTEx fibroblasts

We fitted the pairwise hierarchical model between our eQTLs and the GTEx fibroblast eQTLs^35^ to compute the colocalisation probability. The hypothesis testing was performed with

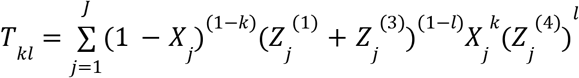

for *k*, *l* = 0, 1, where *X*_*j*_ denotes the probability that the gene *j* is differentially expressed (defined above), *Z*_*j*_^(1)^ denotes the probability that the gene *j* is an eQTL in our data and not in GTEx fibroblasts, *Z*_*j*_^(3)^ denotes the probability that the gene *j* is eQTLs in our data and GTEx fibroblasts and not sharing the putative causal variant, and *Z*_*j*_^(4)^ denotes the probability that the gene j is eQTLs in our data and GTEx fibroblasts and colocalised.

### Colocalisation with GWAS traits

We used the same pairwise hierarchical model^35^ to perform the GWAS colocalisation, where the prior probabilities of the hierarchical model were fixed as {Π_1_, Π_2_, Ψ_3_} = {0. 2, 0. 05, 0. 01}, so that we can compare different studies with different statistical power to detect GWAS associations due to varying sample sizes.

### Annotating TATA and CpG genes

We used the TATA motifs from CIS-BP (Data Availability) and the CpG annotation from UCSC (Data Availability) to annotate genes that have a TATA motif and/or a CpG site 100bp upstream from TSS (we referred to as TATA genes and CpG genes).

### eQTL variant enrichment at transcription factor motifs

The hierarchical model provided the posterior probability that each variant *l* in the cis regulatory region for the gene *j*is the eQTL *Z*_*jl*_ so that 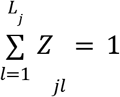 where *L*_*j*_ is the number of variants in the cis window. We first selected the lead eQTL variant according to the posterior probability for each gene *j*. We then used the position weight matrices of transcription factor motifs in CIS-BP (Data Availability) to call motifs overlapping with lead eQTL variants as described elsewhere^34^.

To perform the hypothesis testing that a TF motif is significantly overlapping with eQTL variants, we set 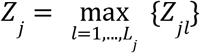 and *X*_*j*_ to be the binary variable whose value is *X*_*j*_ = 1 if the lead eQTL variant *l* is overlapping with a TF motif; otherwise *X*_*j*_ = 0. We then computed the 2×2 table to perform the enrichment analysis as described above.

### PBMC data analysis and eQTL mapping

We reduced the full GASPACHO approach to accommodate the PBMC single cell data of over 700K cells in a reasonable time scale. The kernel functions used in the model were restricted to the linear kernel without the cyclic kernel for the cell cycle effect. The latent factors were estimated with the covariates of the number of genes expressed, the number of mapped reads, the sequencing center, sex, age, COVID19 status, COVID19 sevierity, patient ID, and the first 3 genotype principal components. The latent factors were then used to define the two GPs

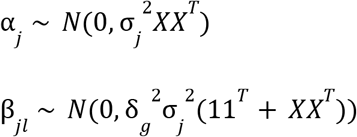

for the intercept and the eQTL effect size of variant *l* for gene *j*.

## GWAS summary statistics

GWAS summary statistics were obtained from Open Targets including studies from GWAS Catalog, FINNGEN and UK Biobank (in total 4,744 traits). The summary statistics were collected and harmonised as described in https://github.com/opentargets/genetics-sumstat-data.

## Data Availability

The annotation of CpG site was downloaded from the UCSC website (https://hgdownload.soe.ucsc.edu/goldenPath/hg38/database/cpgIslandExt.txt.gz). The position weight matrices of transcription factor motifs were obtained from CIS-BP (http://cisbp.ccbr.utoronto.ca/).

---

**Extended Data Figure 1.**
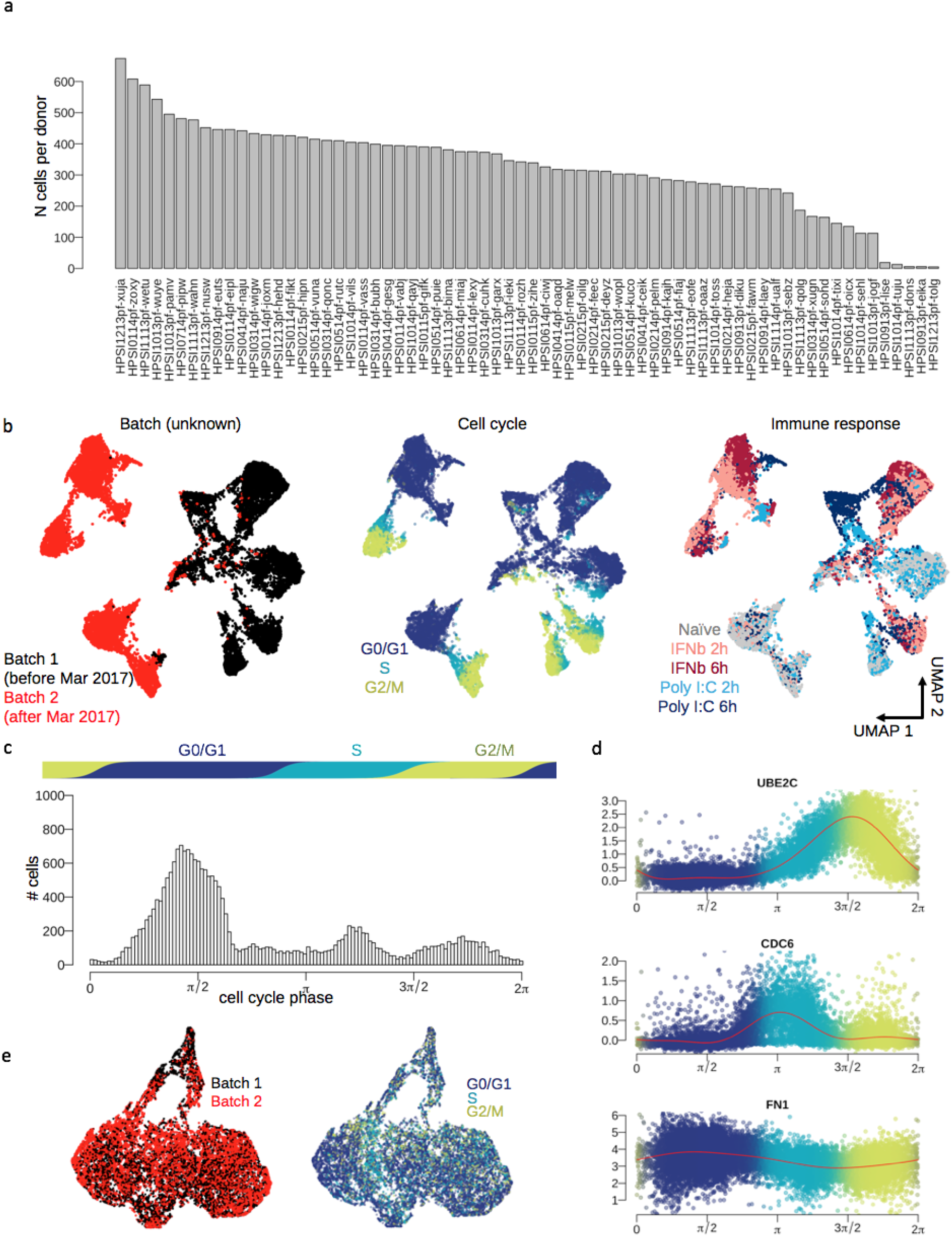
Data and QC. **a.** Barplot shows the number of cells for each donor (cell line). **b.** UMAP calculated from the first XXX principal components from the data. Points are coloured by unknown batch effect (well correlated with experimental date), cell cycle phase estimated from known marker genes (Methods) and experimental conditions. **c.** Histogram shows the distribution of estimated cell cycle phase by GPLVM (Methods). **d.** Scatterplots show scaled expression of known cell cycle genes (UBE2C and CDC6) and a gene highly expressed in G0/G1 phase (FN1). The red curves show the posterior mean estimates of expression levels by GPLVM (Methods). **e.** UMAPs of the target cell states coloured by unknown batch or cell cycle phase.

**Extended Data Figure 2.**
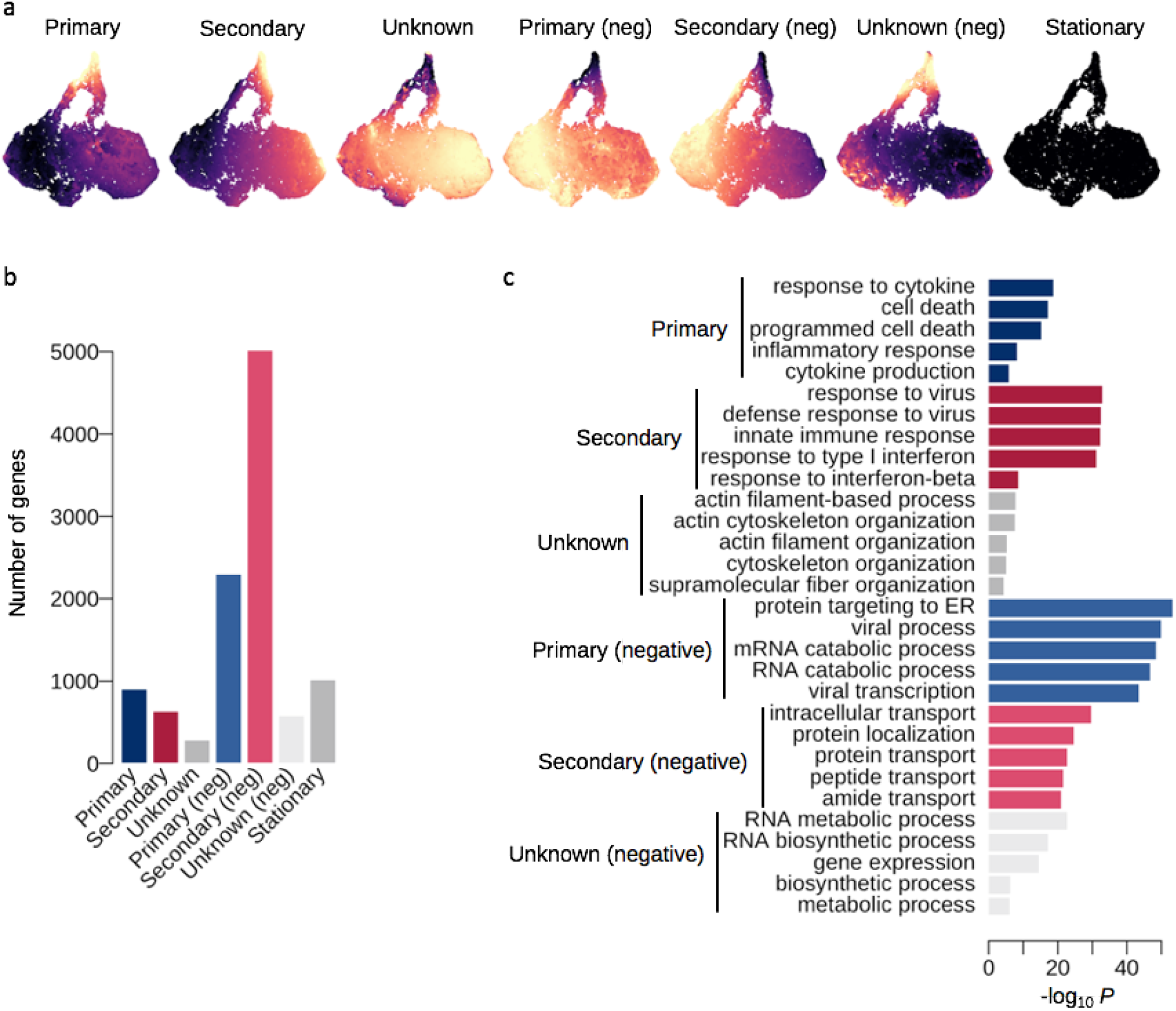
Innate immunity in single cell data. Differential expression result from GPMM. **a**. Seven different mixture components estimated from the GP mixture model (Methods). **b**. The number of genes categorized in each of the seven components. **c**. Top GO terms enriched with the genes detected in each of the seven DE categories.

**Extended Data Figure 3.**
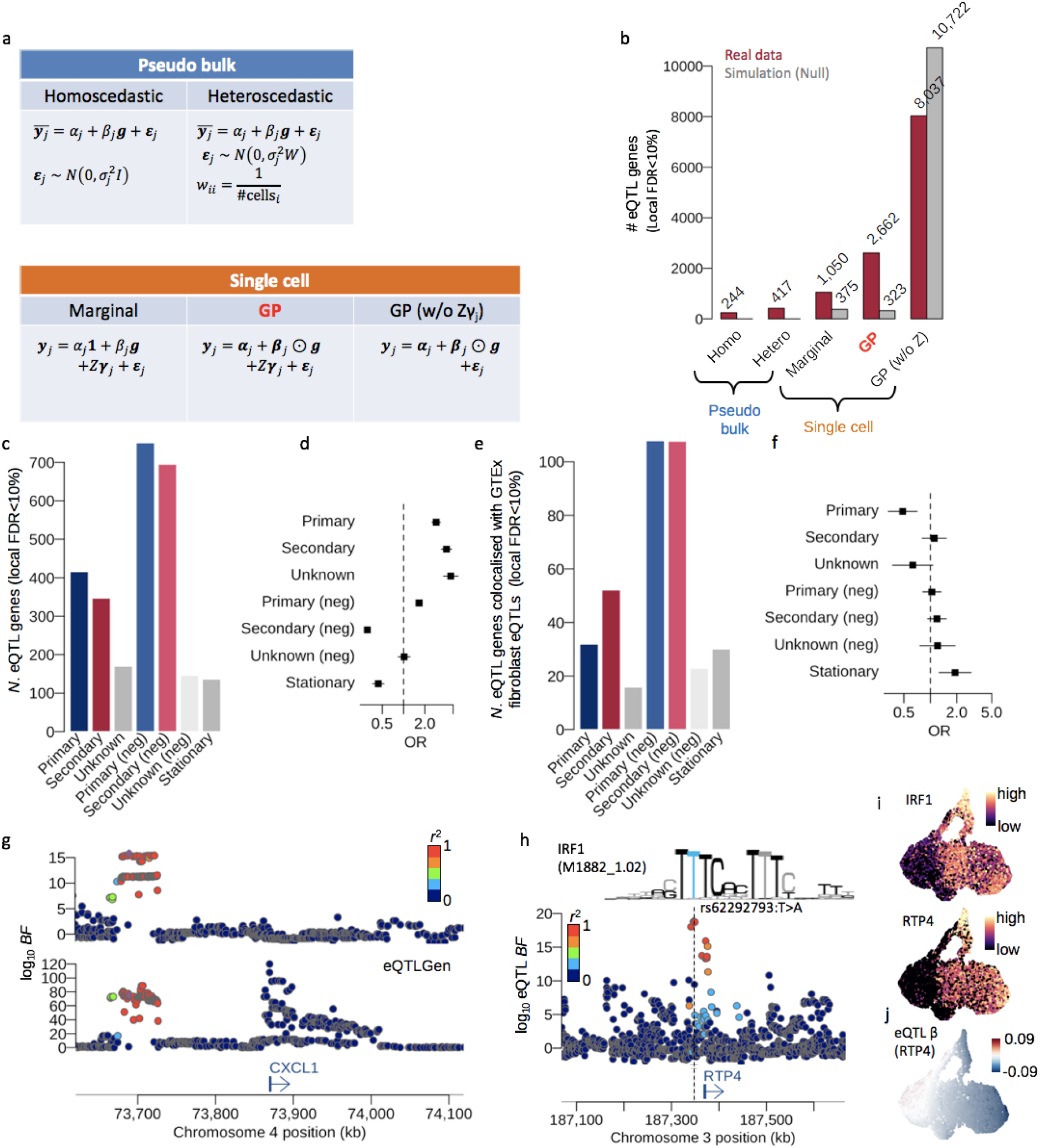
Response eQTL mapping. **a.** Table summarises the 5 different models to map eQTLs (the top two are pseudo bulk based approaches and the bottom three are single cell based approaches). **b.** The numbers of eQTLs mapped by the 5 different models at local FDR 10%. The red bars represent the numbers of eQTL genes for the 5 different methods using the fibroblast data and the gray bars represent the numbers of eQTL genes using a simulated expression data under the null hypothesis (Online Methods). **c.** The number of eQTL genes stratified by the spatial DE genes demonstrated in Extended Data Figure 2b. **d.** Forest plot shows the enrichment of the 2,662 eQTL genes in each DE gene. **e.** The number of eQTL genes colocalised with GTEx fibroblast eQTLs in the 7 different DE gene categories. **f.** Forest plot shows the enrichment of the eQTL colocalised with GTEx fibroblast eQTLs in the 7 different DE gene categories. **g.** Locus zoom plot shows the association Bayes factors of CXCL1 eQTL for our fibroblast (top) and the eQTLGen blood samples (bottom). **h.** Locus zoom plot shows the RTP4 eQTL association Bayes factors. The lead eQTL variant rs62292793T>A disrupts the putative IRF1 binding motif (M1882_1.02; CIS-BP version 1.02) upstream of RTP4 gene promoter. **i.** UMAPs show the expression levels of IRF1 and RTP4. **j.** UMAP shows the eQTL effect size of RTP4.

**Extended Data Figure 4.**
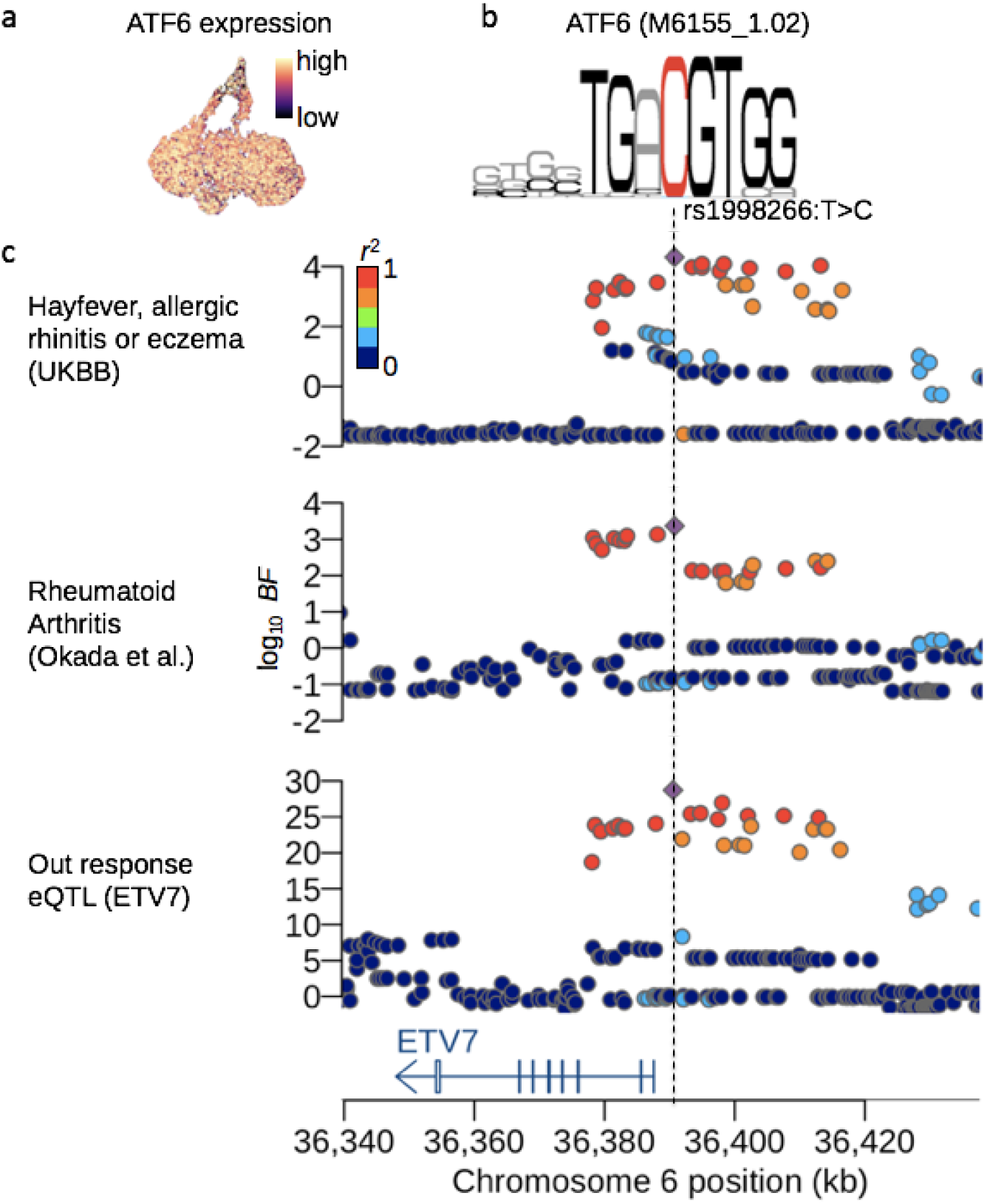
Fine-mapping of ETV7 eQTL. **a.** ATF6 expression on the UMAP. **b.** ATF motif (M6155_1.02; CIS-BP version 1.02). The nucleotide C coloured by red indicates the location of the eQTL variant rs1998266T>C. **c.** Locus zoom plots of hayfever, allergic rhinitis or eczema, rheumatoid arthritis and the ETV7 eQTL.

**Extended Data Figure 5.**
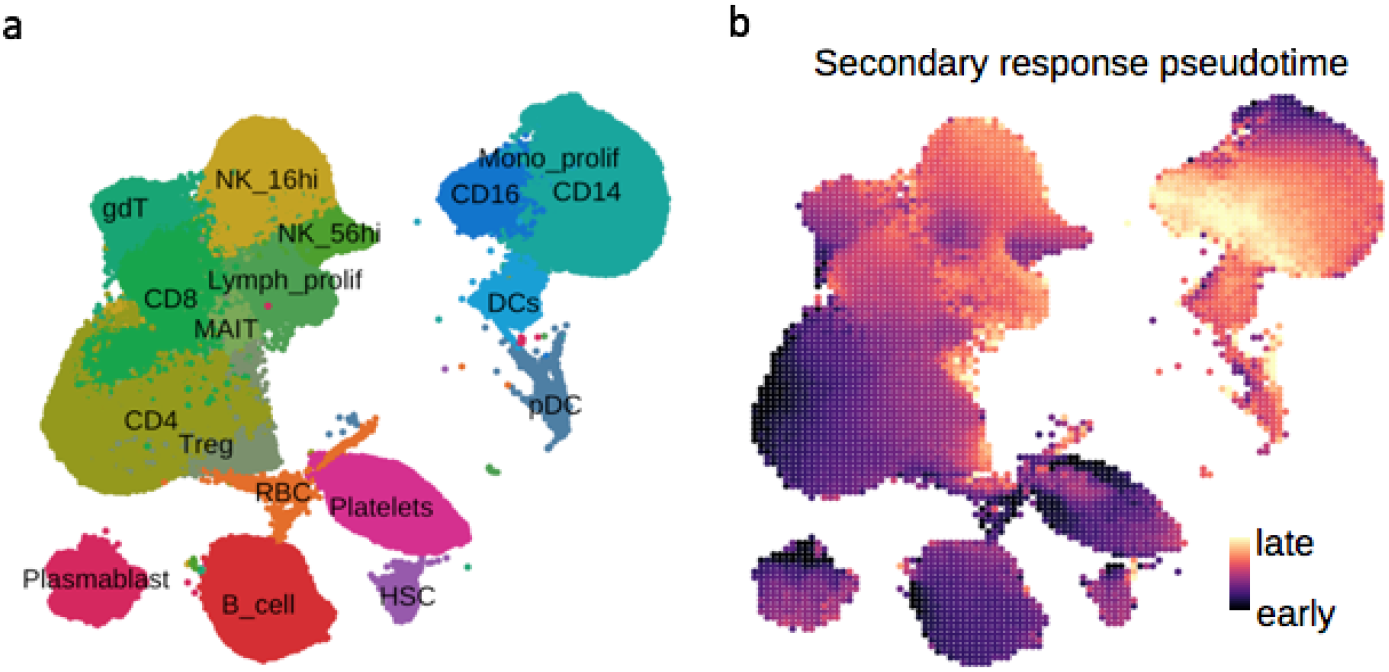
PBMC single cell data. **a.** UMAP shows the 18 different cell types annotated previously. **b.** UMAP shows the secondary response pseudotime calculated from the secondary response genes discovered in the fibroblast data.

**Extended Data Figure 6.**
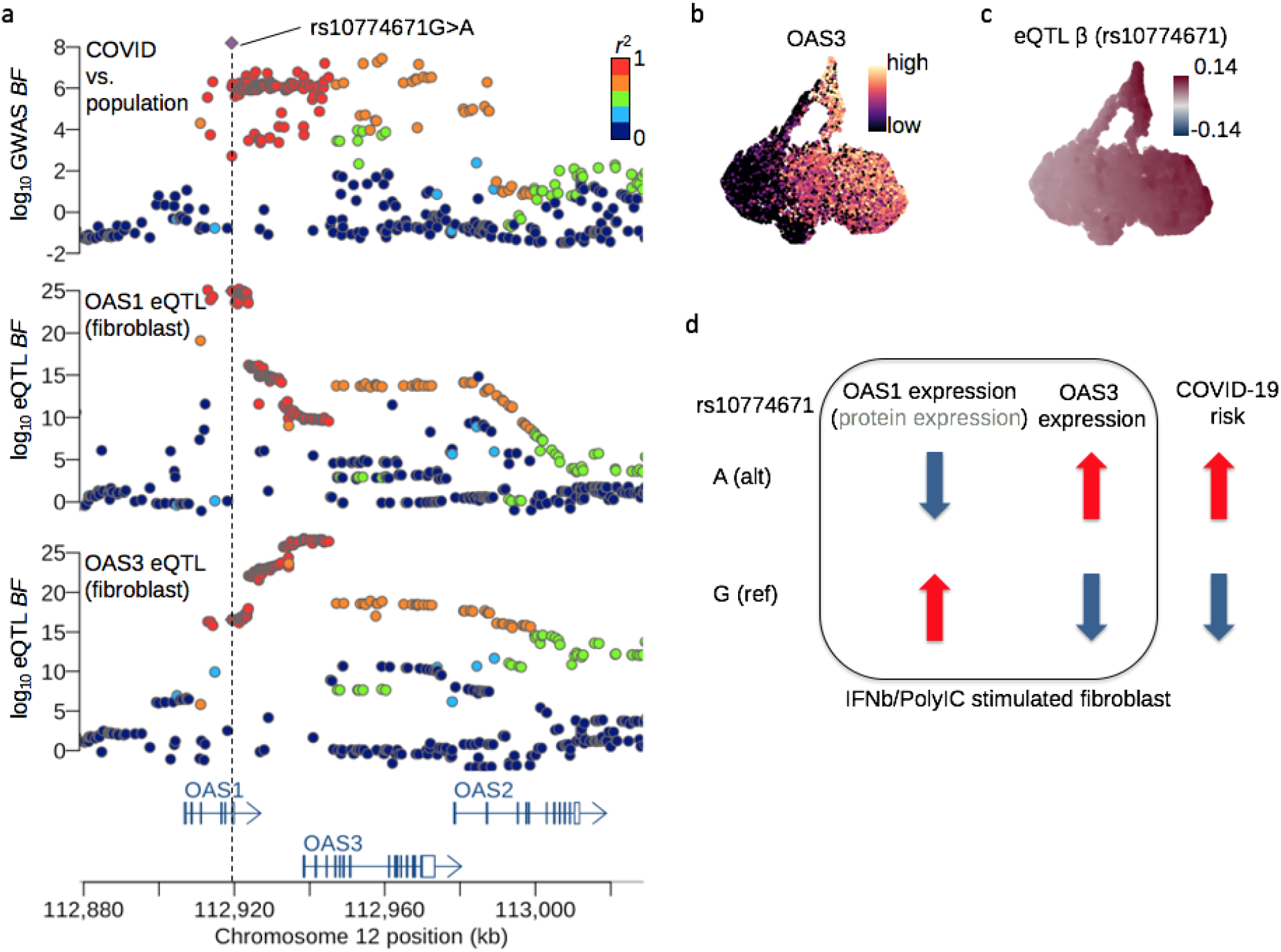
Association and effect directions of OAS3 eQTL in fibroblasts and colocalisation with COVID-19 GWAS. **a.** UMAP shows the OAS3 gene expression. **b.** UMAP shows the eQTL effect size at rs10774671. **c.** Locus zoom plot shows the COVID-19 GWAS, OAS1 and OAS3 eQTL (both in fibroblasts) associations. **d.** Effect direction of OAS1/3 eQTLs and the risk allele of COVID-19 GWAS at the lead variant rs10774671.

**Extended Data Figure 7.**
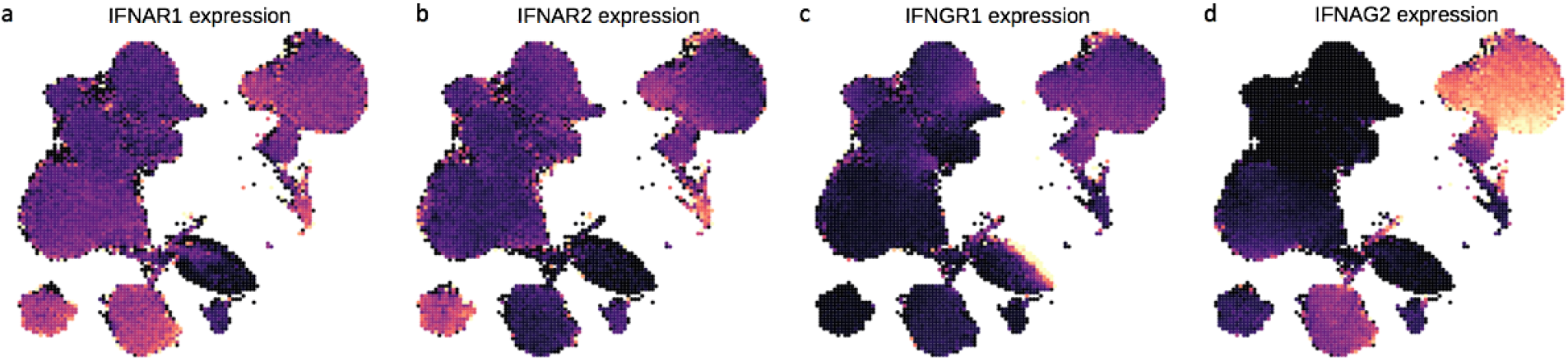
Interferon receptor gene expression. **a.** UMAP shows interferon alpha receptor 1 (IFNAR1) gene expression. **b.** UMAP shows interferon alpha receptor 2 (IFNAR2) gene expression. **c.** UMAP shows interferon gamma receptor 1 (IFNGR1) gene expression. **d.** UMAP shows interferon gamma receptor 1 (IFNGR1) gene expression.

