## Supplementary Note for "Mapping interindividual dynamics of innate immune response at single-cell resolution"

### 1 GASPACHO

The GAuSsian Processes for Association mapping leveraging Cell HeterOgeneity (GASPACHO) is an analysis framework utilising Gaussian Processes (GP) for dimensional reduction and genetic association mapping. GASPACHO consists of three different components: (1) Gaussian Process Latent Variable Model (GPLVM), (2) Gaussian Process mixture model for spatial differential expression anlaysis; and (3) Gaussian Process regression for genetic association mapping in single cell resolution. The subsequent sections demonstrate the model derivation and detailed parameter inference in an ordinary maximum likelihood framework.

#### 1.1 Nomenclature

$N$ : number of cells (samples)
$J$ : number of genes (features)
$M$ : number of inducing points
$O$ : number of fixed variables
$P$ : dimension of design matrix of random effects
$Q$ : number of latent variables
$\mathcal{Y} \in \mathbb{R}^{N\times J}$ : normalised expression matrix
$\mathcal{B} \in \mathbb{R}^{P\times J}$ : random effects
$\mathcal{F} \in \mathbb{R}^{N\times J}$ : sparse GP
$\mathcal{U} \in \mathbb{R}^{M\times J}$ : inducing variables
$\mathcal{X} \in \mathbb{R}^{N\times Q}$ : latent variables ($N$ training points)
$W \in \mathbb{R}^{N\times O}$ : design matrix for fixed effects
$\mathcal{A} \in \mathbb{R}^{N\times O}$ : Fixed effect parameter matrix
$\mathcal{T} \in \mathbb{R}^{M\times Q}$ : inducing points
$Z \in \mathbb{R}^{N\times P}$ : design matrix for random effects
$\zeta \in \mathbb{R}^{P}$ : mean of random effects
$\Delta \in \mathbb{R}^{P\times P}$ : covariance matrix of random effects (diagonal)
$\rho \in \mathbb{R}^{P}$ : length parameters of kernel
$\theta \in \mathbb{R}^{1}$ : variance parameter of kernel
$\Sigma \in \mathbb{R}^{J\times J}$ : residual variance matrix for genes (diagonal)
$\Omega \in \mathbb{R}^{N\times N}$ : residual variance matrix for cells (diagonal)
$\theta K \in \mathbb{R}^{M\times M}$ : Covariance matrix of $M$ inducing points (kernel is defined later)
$\theta \mathcal{K}_{NM} \in \mathbb{R}^{N\times M}$ : Covariance matrix between $N$ training points and $M$ inducing points
$\theta \mathcal{K}_{NN} \in \mathbb{R}^{N\times N}$ : Covariance matrix of $N$ training points
$1_J \in \mathbb{R}^{J}$ : $J$ dimensional vector of all 1's
$I_N \in \mathbb{R}^{N\times N}$ : $N$ dimensional identity matrix
$\mathcal{N}(Y|M,U,V)$ : matrix normal distribution of random variable matrix $Y \in \mathbb{R}^{N\times P}$ with mean $M \in \mathbb{R}^{N\times P}$, row variance matrix $U \in \mathbb{R}^{N\times N}$ and column variance matrix $V \in \mathbb{R}^{P\times P}$

#### 1.2 Kernel

We assume gene expression is affected by cell cycle and other biological and technical effects. Those effects are not known a priori, and therefore to be inferred from the data. For $Q$-dimensional latent vectors $x, x' \in \mathbb{R}^Q$, the kernel function specifically used is given by the product of the periodic and ARD SE kernel:

$$\theta k(x, x') = \theta \exp\left\{-\frac{2\sin^2(|x_1 - x'_1|/2)}{\rho_1^2}\right\} \exp\left\{-\sum_{q=2}^{Q} \frac{(x_q - x'_q)^2}{2\rho_q^2}\right\},$$

where the periodic term models the cell cycle effect and the ARD SE kernel for the other effects.

For the $N$ training points $\mathcal{X}^\top = (x_k \in \mathbb{R}^Q; 1 \leq k \leq N)$ and $M$ inducing points $\mathcal{T}^\top = (t_k \in \mathbb{R}^Q; 1 \leq k \leq M)$, we define the covariance matrices:

$$\begin{aligned} K &= \{k(t_k, t_l); 1 \leq k, l \leq M\}, \\ \mathcal{K}_{NM} &= \{k(x_k, t_l); 1 \leq k \leq N, 1 \leq l \leq M\}, \\ \mathcal{K}_{NN} &= \{k(x_k, x_l); 1 \leq k, l \leq N\}. \end{aligned}$$

#### 1.3 GPLVM

Our GPLVM is an extension of Titsias and Lawrence [] with introducing both fixed and random effect terms that internally adjust the expression matrix $\mathcal{Y}$. The joint probability of the GPLVM is written as a product of matrix normal distributions:

$$\begin{aligned} p(\mathcal{Y}, \mathcal{B}, \mathcal{F}, \mathcal{U}, \mathcal{X}) = \mathcal{N}(\mathcal{Y}|W\mathcal{A} + Z\mathcal{B} + \mathcal{F}, \Omega, \Sigma)\mathcal{N}(\mathcal{B}|\zeta 1_J^\top, \Delta, \Sigma) \\ \times \mathcal{N}(\mathcal{F}|\mathcal{K}_{NM}K^{-1}\mathcal{U}, \theta\tilde{\mathcal{K}}_{NN}, \Sigma)\mathcal{N}(\mathcal{U}|0, \theta K, \Sigma)\mathcal{N}(\mathcal{X}|0, I_N, I_Q), \end{aligned}$$

where $\tilde{\mathcal{K}}_{NN} = \mathcal{K}_{NN} - \mathcal{K}_{NM}K^{-1}\mathcal{K}_{MN}$. Because $\mathcal{K}_{NN}$ is not tractable for large $N$, we calculate the lower bound

$$\begin{aligned} \log p(\mathcal{Y}|\mathcal{B}, \mathcal{U}, \mathcal{X}) &= \log \int p(\mathcal{F}|\mathcal{U}) p(\mathcal{Y}|\mathcal{B}, \mathcal{F}, \mathcal{U}) d\mathcal{F} \\ &\geq \int p(\mathcal{F}|\mathcal{U}) \log p(\mathcal{Y}|\mathcal{B}, \mathcal{F}, \mathcal{U}) d\mathcal{F} \\ &= \log p(\mathcal{Y}|W\mathcal{A} + Z\mathcal{B} + \mathcal{K}_{NM}K^{-1}\mathcal{U}, \Omega, \Sigma) - \frac{J}{2}\mathrm{tr}(\Omega^{-1}\theta\tilde{\mathcal{K}}_{NN}) \\ &\equiv \mathcal{L}_1. \end{aligned}$$

We then have the marginal probability of $\mathcal{Y}$ given $\mathcal{X}$ as the Titsias lower bound, such that

$$\begin{aligned} p(\mathcal{Y}|\mathcal{X})p(\mathcal{X}) &\geq p(\mathcal{X}) \int \exp\{\mathcal{L}_1\} p(\mathcal{B}) p(\mathcal{U}) d\mathcal{B} d\mathcal{U} \\ &= \mathcal{N}(\mathcal{Y}|W\mathcal{A} + Z\zeta 1_J^\top, \Omega + Z\Delta Z^\top + \theta\mathcal{K}_{NM}K^{-1}\mathcal{K}_{MN}, \Sigma) \\ &\quad \times \mathcal{N}(\mathcal{X}|0, I_N, I_Q) \exp\{-J\mathrm{tr}(\Omega^{-1}\theta\tilde{\mathcal{K}}_{NN})/2\} \\ &\equiv \exp\{\mathcal{L}_2\}. \end{aligned}$$

We maximise $\mathcal{L}_2$ with respect to $\Theta \equiv \{\Sigma, \Omega, \mathcal{X}, \mathcal{T}, \rho, \theta, \Delta, \zeta, \mathcal{A}\}$ using the quasi Newtom method (L-BFGS) with the gradient analytically obtained.

After some matrix algebra, the Titsias bound of our model can be written as

$$\mathcal{L}_2 = -\frac{NJ}{2}\log(2\pi) - \frac{N}{2}\log|\Sigma| - \frac{J}{2}\log|V| - \frac{J}{2}\mathrm{tr}(SV^{-1}) - \frac{J}{2}\mathrm{tr}(\Omega^{-1}\theta\tilde{\mathcal{K}}_{NN}),$$

where

$$\begin{aligned}
V &= \Omega + \tilde{Z}\tilde{\Delta}\tilde{Z}^\top, \\
V^{-1} &= \Omega^{-1} - \Omega^{-1}\tilde{Z}\Phi\tilde{Z}^\top\Omega^{-1}, \\
S &= (\mathcal{Y} - \tilde{W}\tilde{\mathcal{A}})\Sigma^{-1}(\mathcal{Y} - \tilde{W}\tilde{\mathcal{A}})^\top / J
\end{aligned}$$

with modified matrix notations

$$\begin{aligned}
\tilde{Z} &= (Z, \mathcal{K}_{NM}) \\
\tilde{\Delta} &= \begin{pmatrix} \Delta & 0 \\ 0 & \theta K^{-1} \end{pmatrix} \\
\tilde{W} &= (W, Z\zeta) \\
\tilde{\mathcal{A}} &= \begin{pmatrix} \mathcal{A} \\ 1_J^\top \end{pmatrix} \\
\Phi^{-1} &= \tilde{\Delta}^{-1} + \tilde{Z}^\top\Omega^{-1}\tilde{Z}
\end{aligned}$$

to make the lower bound simpler.

The model parameters are alternatively updated to maximise the lower bound where some of the model parameters have the analytical solution at each iteration, while others don't. The following subsections provide either the first derivative of the lower bound (the gradient) or the exact solution of each model parameters where the gradient to be 0.

###### 1.3.1 The fixed parameters $\mathcal{A}$ and $\eta$

The fixed parameters have the exact solution at each iteration of optimisation. The maximum of the lower bound is attained by

$$\begin{aligned}
\hat{\zeta} &= (Z^\top V^{-1} Z + \lambda I_P)^{-1} Z^\top V^{-1}[(1_N^\top \Sigma^{-1}\mathcal{Y}^\top)^\top - W\mathcal{A}\Sigma^{-1}1_N], \\
\hat{\mathcal{A}} &= (W^\top V^{-1} W)^{-1} W^\top V^{-1}(\mathcal{Y} - Z\zeta 1_J^\top),
\end{aligned}$$

where $\lambda > 0$ is a ridge parameter to avoid the parameter undetermined, because $\mathrm{rank}(Z) < P$ in general. We set $\lambda = 0.01$ in the current implementation. Note that, when we optimise $\mathcal{L}_2$ with respect to $\zeta$ or $\mathcal{A}$, all there parameters are fixed constant at each iteration.

###### 1.3.2 Gene specific residual variance $\Sigma$

The matrix derivative of $\mathcal{L}_2$ with respect to the precision $\Sigma^{-1}$ (but not variance $\Sigma$) gives

$$\partial_{\Sigma^{-1}}\mathcal{L}_2 = \frac{N}{2}\mathrm{tr}(\Sigma\partial\Sigma^{-1}) - \frac{1}{2}\mathrm{tr}\left[\partial\Sigma^{-1}(\mathcal{Y} - \tilde{W}\tilde{\mathcal{A}})^\top V^{-1}(\mathcal{Y} - \tilde{W}\tilde{\mathcal{A}})\right],$$

suggesting the diagonal element of $(\mathcal{Y} - \tilde{W}\tilde{\mathcal{A}})^\top V^{-1}(\mathcal{Y} - \tilde{W}\tilde{\mathcal{A}})/N$ gives the optimum variance parameter. Note that it requires the matrix computation

$$\begin{aligned}
(\mathcal{Y} - \tilde{W}\tilde{\mathcal{A}})^\top V^{-1}(\mathcal{Y} - \tilde{W}\tilde{\mathcal{A}}) = {}& \mathcal{Y}^\top\Omega^{-1}\mathcal{Y} - \mathbb{C}\tilde{\mathcal{A}} - \tilde{\mathcal{A}}^\top\mathbb{C}^\top + \tilde{\mathcal{A}}^\top(\tilde{W}^\top\Omega^{-1}\tilde{W})\tilde{\mathcal{A}} \\
& - \mathbb{D}\Phi\mathbb{D}^\top + \mathbb{D}\Phi\mathbb{E}\tilde{\mathcal{A}} + \tilde{\mathcal{A}}^\top\mathbb{E}^\top\Phi\mathbb{D}^\top - \tilde{\mathcal{A}}^\top\mathbb{E}^\top\Phi\mathbb{E}\tilde{\mathcal{A}}
\end{aligned}$$

where

$$\begin{aligned}
\mathbb{C} &\equiv \mathcal{Y}^\top\Omega^{-1}\tilde{W}, \\
\mathbb{D} &\equiv \mathcal{Y}^\top\Omega^{-1}\tilde{Z}, \\
\mathbb{E} &\equiv \tilde{Z}^\top\Omega^{-1}\tilde{W}
\end{aligned}$$

should be precomputed for a scalable optimisation.

Although the variance parameter in the original model above has the exact solution, the model can be easily extended when the parameter estimate is unstable because of many 0's in the expression data. Then we can assume the precision parameter to be Gamma distributed with the following shape and rate parameters:

$$\frac{1}{\sigma_j^2} \equiv \tau_j \sim \mathcal{G}(\nu/2, \nu\mu_j/2)$$
$$\mu_j = x_j^\top \lambda$$

so that $\mathbb{E}[1/\tau_j] = \mathbb{E}[\sigma_j^2] = \mu_j$ and $x_j$ is an arbitrary covariate vector. We normally use the percentage $p_j$ of cells that express each gene $j$ and $x_j^\top = (p_j, p_j - p_j^2)$. Here the Gamma distribution is the conjugate prior on $\mathcal{L}_2$, we can analytically obtain the posterior probability distribution of $\tau_j$:

$$\tau_j|\mathcal{Y} \sim \mathcal{G}\left(\frac{\nu+N}{2}, \frac{\nu\mu_j + SSE_j}{2}\right),$$

where $SSE_j$ is the $j$th diagonal element of $(\mathcal{Y} - \tilde{W}\tilde{\mathcal{A}})^\top V^{-1}(\mathcal{Y} - \tilde{W}\tilde{\mathcal{A}})$. The hyperparameter $\lambda$ can be learned from the data in a standard generalised linear model with Gamma outcome, where the dependent variable $\tau_j$ and its $\log \tau_j$ are replaced at each iteration by the posterior means

$$\mathbb{E}[\tau_j|\mathcal{Y}] = \frac{\nu+N}{\nu\mu_j + SSE_j},$$
$$\mathbb{E}[\log \tau_j|\mathcal{Y}] = \psi\left(\frac{\nu+N}{2}\right) - \log\left(\frac{\nu\mu_j + SSE_j}{2}\right),$$

where$\psi(\cdot)$ is the digamma function.

##### 1.3.3 Cell specific residual variance $\Omega$

Because we introduce the column variance $\Sigma$ and the row variance $\Omega$ for $\mathcal{Y}$, there is a parameter uncertainty. To overcome this issue and guarantee the uniqueness of parameter estimation, we assume an inverse Gamma distribution with fixed shape and rate parameters ($\alpha$ and $\beta$ respectively) on each cell specific residual variance for cell $i$ (the $i$th diagonal element of $\Omega$), such that

$$\omega_i^{-2} \sim \mathcal{G}(\alpha, \beta).$$

This implies that we maximise $\tilde{\mathcal{L}}_2 \equiv \mathcal{L}_2 + (\alpha - 1)\log|\Omega^{-1}| - \beta \mathrm{tr}(\Omega^{-1})$ instead of $\mathcal{L}_2$ for $\Omega$. The first derivative with respect to $\Omega^{-1}$ is given by

$$\partial_{\Omega^{-1}}\tilde{\mathcal{L}}_2 = \frac{J}{2}\mathrm{tr}\{\Omega V^{-1}\Omega(\partial\Omega^{-1})\} - \frac{J}{2}\mathrm{tr}\{\Omega V^{-1} S V^{-1}\Omega(\partial\Omega^{-1})\} - \frac{J}{2}\mathrm{tr}\{\theta\tilde{\mathcal{K}}_{NN}(\partial\Omega^{-1})\} + (\alpha-1)\mathrm{tr}\{\Omega(\partial\Omega^{-1})\} - \beta\mathrm{tr}\{\partial\Omega^{-1}\}.$$

In order to make a scalable computation, we calculate

$$\Omega V^{-1} S V^{-1}\Omega = (\mathcal{Y}\Sigma^{-1}\mathcal{Y}^\top - \mathcal{Y}\Sigma^{-1}\mathbb{D}\Phi\tilde{Z}^\top + \mathcal{Y}\Sigma^{-1}\tilde{\mathcal{A}}^\top\mathbb{F}^\top - \tilde{Z}\Phi\mathbb{D}^\top\Sigma^{-1}\mathcal{Y}^\top + \tilde{Z}\Phi\mathbb{D}^\top\Sigma^{-1}\mathbb{D}\Phi\tilde{Z}^\top - \tilde{Z}\Phi\mathbb{D}^\top\Sigma^{-1}\tilde{\mathcal{A}}^\top\mathbb{F}^\top + \mathbb{F}\tilde{\mathcal{A}}\Sigma^{-1}\mathcal{Y}^\top - \mathbb{F}\tilde{\mathcal{A}}\Sigma^{-1}\mathbb{D}\Phi\tilde{Z}^\top + \mathbb{F}\tilde{\mathcal{A}}\Sigma^{-1}\tilde{\mathcal{A}}^\top\mathbb{F}^\top)/J$$

with precomputed matrices

$$\mathbb{D} = \mathcal{Y}^\top\Omega^{-1}\tilde{Z},$$
$$\mathbb{E} = \tilde{Z}^\top\Omega^{-1}\tilde{W},$$
$$\mathbb{F} = -\tilde{W} + \tilde{Z}\Phi\mathbb{E}.$$

##### 1.3.4 Variance parameters $\tilde{\Delta}$ for the random effects and GP

There is no closed form for the solution of $\tilde{\Delta}$, therefore the first derivative

$$\begin{aligned}\partial_{\tilde{\Delta}}\mathcal{L}_2 &= -\frac{J}{2}\text{tr}\{V^{-1}(\partial_{\tilde{\Delta}}V)\} + \frac{J}{2}\text{tr}\{V^{-1}SV^{-1}(\partial_{\tilde{\Delta}}V)\} - \frac{J}{2}\text{tr}\{(\partial\theta)\Omega^{-1}(I - \mathcal{K}_{NM}K^{-1}\mathcal{K}_{MN})\} \\ &= -J\text{tr}\{\tilde{\Delta}\tilde{Z}^\top V^{-1}(\partial\tilde{Z})\} + J\text{tr}\{\tilde{\Delta}\tilde{Z}^\top V^{-1}SV^{-1}(\partial\tilde{Z})\} \\ &\quad - \frac{J}{2}\text{tr}\{\tilde{Z}^\top V^{-1}\tilde{Z}(\partial\tilde{\Delta})\} + \frac{J}{2}\text{tr}\{\tilde{Z}^\top V^{-1}SV^{-1}\tilde{Z}(\partial\tilde{\Delta})\} - \frac{J}{2}\text{tr}\{(\partial\theta)\Omega^{-1}(I - \mathcal{K}_{NM}K^{-1}\mathcal{K}_{MN})\}\end{aligned}$$

is used to update $\tilde{\Delta}$ by using L-BFGS algorithm, where

$$\begin{aligned}\tilde{Z}^\top V^{-1} &= \tilde{\Delta}^{-1}\Phi\tilde{Z}^\top\Omega^{-1}, \\ \tilde{Z}^\top V^{-1}\tilde{Z} &= \tilde{\Delta}^{-1}(\tilde{\Delta} - \Phi)\tilde{\Delta}^{-1}.\end{aligned}$$

After the matrix manipulation, we obtain

$$\partial_V\mathcal{L}_2 = -\frac{J}{2}\text{tr}\{\mathbb{A}(\partial\tilde{\Delta})\} + \frac{J}{2}\text{tr}\{\mathbb{B}(\partial\tilde{\Delta})\} - \frac{J}{2}\text{tr}\{\Omega^{-1}(I - \mathcal{K}_{NM}K^{-1}\mathcal{K}_{MN})\}(\partial\theta),$$

where

$$\begin{aligned}\mathbb{A} &\equiv \tilde{Z}^\top V^{-1}\tilde{Z} = \tilde{\Delta}^{-1}(\tilde{\Delta} - \Phi)\tilde{\Delta}^{-1}, \\ \mathbb{B} &\equiv \tilde{\Delta}^{-1}\Phi\tilde{Z}^\top\Omega^{-1}S\Omega^{-1}\tilde{Z}\Phi\tilde{\Delta}^{-1} = \tilde{\Delta}^{-1}\mathbb{C}\Sigma^{-1}\mathbb{C}^\top\tilde{\Delta}^{-1}/J, \\ \mathbb{C} &\equiv \Phi(\mathbb{D}^\top - \mathbb{E}\tilde{\mathcal{A}}), \\ \mathbb{D} &\equiv \mathcal{Y}^\top\Omega^{-1}\tilde{Z}, \\ \mathbb{E} &\equiv \tilde{Z}^\top\Omega^{-1}\tilde{W}\end{aligned}$$

should again be precomputed for a scalable optimisation.

##### 1.3.5 Latent variables $\mathcal{X}$ and $\mathcal{T}$

There is also no closed form for the solution of $\mathcal{X}$, we here provide the first derivative

$$\begin{aligned}\partial_{\mathcal{X}}\mathcal{L}_2 &= -\frac{J}{2}\text{tr}\{V^{-1}(\partial_{\mathcal{X}}V)\} + \frac{J}{2}\text{tr}\{V^{-1}SV^{-1}(\partial_{\mathcal{X}}V)\} - \frac{J}{2}\text{tr}\{\Omega^{-1}\theta\partial_{\mathcal{X}}(I - \mathcal{K}_{NM}K^{-1}\mathcal{K}_{MN})\} \\ &= -J\text{tr}\{\tilde{\Delta}\tilde{Z}^\top V^{-1}(\partial_{\mathcal{X}}\tilde{Z})\} + J\text{tr}\{\tilde{\Delta}\tilde{Z}^\top V^{-1}SV^{-1}(\partial_{\mathcal{X}}\tilde{Z})\} + \frac{J}{2}\text{tr}\{\Omega^{-1}\theta\partial_{\mathcal{X}}(\mathcal{K}_{NM}K^{-1}\mathcal{K}_{MN})\} \\ &= -J\text{tr}\{\Phi\tilde{Z}^\top\Omega^{-1}(0, \partial_{\mathcal{X}}\mathcal{K}_{NM})\} + J\text{tr}\{\Phi\tilde{Z}^\top\Omega^{-1}SV^{-1}(0, \partial_{\mathcal{X}}\mathcal{K}_{NM})\} + J\text{tr}\{\theta K^{-1}\mathcal{K}_{MN}\Omega^{-1}(\partial_{\mathcal{X}}\mathcal{K}_{NM})\}\end{aligned}$$

with

$$\begin{aligned}\Phi\tilde{Z}^\top\Omega^{-1}SV^{-1} &= \Phi\tilde{Z}^\top\Omega^{-1}(\mathcal{Y} - \tilde{W}\tilde{\mathcal{A}})\Sigma^{-1}(\mathcal{Y}^\top\Omega^{-1} - \mathbb{D}\Phi\tilde{Z}^\top\Omega^{-1} + \tilde{\mathcal{A}}^\top\mathbb{F}^\top\Omega^{-1})/J \\ &= \Phi(\mathbb{D}^\top - \mathbb{E}\tilde{\mathcal{A}})\Sigma^{-1}(\mathcal{Y}^\top\Omega^{-1} - \mathbb{D}\Phi\tilde{Z}^\top\Omega^{-1} + \tilde{\mathcal{A}}^\top\mathbb{F}^\top\Omega^{-1})/J \\ &= \Phi(\mathbb{D}^\top\Sigma^{-1}\mathcal{Y}^\top - \mathbb{E}\mathbb{H} - \mathbb{G}\mathbb{D}\Phi\tilde{Z}^\top + \mathbb{G}\tilde{\mathcal{A}}^\top\mathbb{F}^\top)\Omega^{-1}/J,\end{aligned}$$

where the following matrices are precomputed a priori:

$$\begin{aligned}\mathbb{D} &= \mathcal{Y}^\top\Omega^{-1}\tilde{Z}, \\ \mathbb{E} &= \tilde{Z}^\top\Omega^{-1}\tilde{W}, \\ \mathbb{F} &= -\tilde{W} + \tilde{Z}\Phi\mathbb{E}, \\ \mathbb{G} &= (\mathbb{D}^\top - \mathbb{E}\tilde{\mathcal{A}})\Sigma^{-1}, \\ \mathbb{H} &= \tilde{\mathcal{A}}\Sigma^{-1}\mathcal{Y}^\top.\end{aligned}$$

Likewise,

$$\partial_{\mathcal{T}}\mathcal{L}_2 = -J\mathrm{tr}\{\Phi\tilde{Z}^\top\Omega^{-1}(0,\partial_{\mathcal{T}}\mathcal{K}_{NM})\} + J\mathrm{tr}\{\Phi\tilde{Z}^\top\Omega^{-1}SV^{-1}(0,\partial_{\mathcal{T}}\mathcal{K}_{NM})\} + J\mathrm{tr}\{\theta K^{-1}\mathcal{K}_{MN}\Omega^{-1}(\partial_{\mathcal{T}}\mathcal{K}_{NM})\}.$$

###### 1.3.6 The length parameter $\rho$ in the kernel

Similar calculation is also required for the length parameter of the kernel. The first derivative with respect to $\rho$ is then given by

$$\begin{aligned}\partial_\rho\mathcal{L}_2 = &-J\mathrm{tr}\{\Phi\tilde{Z}^\top\Omega^{-1}(0,\partial\mathcal{K}_{NM})\} + J\mathrm{tr}\{\Phi\tilde{Z}^\top\Omega^{-1}SV^{-1}(0,\partial\mathcal{K}_{NM})\} + J\mathrm{tr}\{\theta K^{-1}\mathcal{K}_{MN}\Omega^{-1}(\partial\mathcal{K}_{NM})\}\\ &+\frac{J}{2}\mathrm{tr}\left\{(\tilde{\Delta}-\Phi)\begin{pmatrix}0 & 0\\ 0 & \theta^{-1}\partial K\end{pmatrix}\right\} - \frac{J}{2}\mathrm{tr}\left\{\Phi\tilde{Z}^\top\Omega^{-1}S\Omega^{-1}\tilde{Z}\Phi\begin{pmatrix}0 & 0\\ 0 & \theta^{-1}\partial K\end{pmatrix}\right\}\\ &-\frac{J}{2}\mathrm{tr}\{\theta K^{-1}\mathcal{K}_{MN}\Omega^{-1}\mathcal{K}_{NM}K^{-1}(\partial K)\}\end{aligned}$$

where

$$\begin{aligned}\Phi\tilde{Z}^\top\Omega^{-1}S\Omega^{-1}\tilde{Z}^\top\Phi &= \Phi(\mathbb{D}^\top - \mathbb{E}\tilde{\mathcal{A}})\Sigma^{-1}(\mathbb{D}^\top - \mathbb{E}\tilde{\mathcal{A}})^\top\Phi/J,\\ \mathbb{D} &= \mathcal{Y}^\top\Omega^{-1}\tilde{Z},\\ \mathbb{E} &= \tilde{Z}^\top\Omega^{-1}\tilde{W}.\end{aligned}$$

#### 1.4 GP mixture model for spatial differential expression

Once we fit the GPLVM and obtained the kernel parameters and latent variables, we then split $\mathcal{X}$ into the target cell state $\mathcal{X}_1$ and the rest $\mathcal{X}_2$ for other technical or unexpected biological variations including cell cycle variation (likewise $\mathcal{T}$ into $\mathcal{T}_1$ and $\mathcal{T}_2$). Then we compute the kernel matrices

$$\begin{aligned}\mathcal{K}_{NN}^{(i)} &= k_i(\mathcal{X}_i,\mathcal{T}_i),\\ \mathcal{K}_{NM}^{(i)} &= k_i(\mathcal{X}_i,\mathcal{T}_i),\\ K^{(i)} &= k_i(\mathcal{T}_i,\mathcal{T}_i),\end{aligned}$$

for $i = 1, 2$, where

$$\begin{aligned}\theta k_1(x,x') &= \theta\exp\left\{-\sum_{q\in Q_1}\frac{(x_q-x'_q)^2}{2\rho_q^2}\right\},\\ \theta k_2(x,x') &= \theta\exp\left\{-\frac{2\sin^2(|x_1-x'_1|/2)}{\rho_1^2}\right\}\exp\left\{-\sum_{q\in Q_2}\frac{(x_q-x'_q)^2}{2\rho_q^2}\right\},\end{aligned}$$

and $Q_1$ denotes the index set of target latent variables and $Q_2$ is the complement set.

Let us denote by $\tilde{y}_j \equiv (y_j - W\hat{a}_j - Z\hat{\zeta})/\hat{\sigma}_j$ the normalised expression for gene $j$. We assume the gene $j$ belongs to one of the $C$ differential expression categories in the target space, in which $f_c$ denotes a GP capturing the $c$th spatial expression pattern ($c = 1, \ldots, C$). Specifically, we model

$$\begin{aligned}\tilde{y}_j|f_c,f_j,\delta_{cj},b_j &\sim \mathcal{N}(\delta_{cj}f_c + f_j + Zb_j,\hat{\Omega})\\ f_c &\sim \mathcal{N}(0,\hat{\theta}\mathcal{K}_{NN}^{(1)})\\ f_j &\sim \mathcal{N}(0,\hat{\theta}\mathcal{K}_{NN}^{(2)})\\ b_j &\sim \mathcal{N}(\hat{\zeta},\hat{\Delta})\\ \delta_{cj} &\sim \mathcal{N}(0,1)\end{aligned}$$

where $\delta_{cj}$ is a coefficient to allow the direction and magnitude of the $c$th spatial component varying across genes. Note that $C = 3$ in our result.

In reality, $\mathcal{K}^{(1)}_{NN}$ or $\mathcal{K}^{(2)}_{NN}$ are not tractable for large $N$, we introduce the inducing variables $\beta_c$ and $u_j$, such that

$$\begin{aligned}
f_c|\beta_c &\sim \mathcal{N}(X\beta_c, \hat{\theta}\tilde{\mathcal{K}}^{(1)}_{NN}),\\
f_j|u_j &\sim \mathcal{N}(\mathcal{K}^{(2)}_{NM}{K^{(2)}}^{-1}u_j, \hat{\theta}\tilde{\mathcal{K}}^{(2)}_{NN}),\\
\beta_c &\sim \mathcal{N}(0, \hat{\theta}K^{(1)}),\\
u_j &\sim \mathcal{N}(0, \hat{\theta}K^{(2)}),
\end{aligned}$$

with $X = \mathcal{K}^{(1)}_{NM}{K^{(1)}}^{-1}$ to compute the lower bound

$$\begin{aligned}
\log p(\tilde{y}_j|\beta_c, u_j, \delta_{cj}, b_j) &= \log \int p(\tilde{y}_j|f_c, f_j, \delta_{cj}, b_j)p(f_c|\beta_c)p(f_j|u_j)df_c df_j\\
&\geq \int \log p(\tilde{y}_j|f_c, f_j, \delta_{jc}, b_j)p(f_c|\beta_c)p(f_j|u_j)df_c df_j\\
&= \log \mathcal{N}(\tilde{y}_j|\delta_{cj}\bar{f}_c + \bar{f}_j + Zb_j, \hat{\Omega}) - \frac{1}{2}\mathrm{tr}\{\hat{\Omega}^{-1}\hat{\theta}(\delta^2_{cj}\tilde{\mathcal{K}}^{(1)}_{NN} + \tilde{\mathcal{K}}^{(2)}_{NN})\}\\
&\equiv \mathcal{L}^{(1)}_{cj}.
\end{aligned}$$

Then Titsias lower bounds can be obtained by

$$\begin{aligned}
p_0(\tilde{y}_j) &\geq \int \exp\{\mathcal{L}^{(1)}_{cj}|_{\beta_c=0,\delta_{cj}=0}\}p(u_j)p(b_j)du_j db_j\\
&= \mathcal{N}(\tilde{y}_j|0, \hat{\Omega} + Z\hat{\Delta}Z^\top + \hat{\theta}\mathcal{K}^{(2)}_{NM}{K^{(2)}}^{-1}\mathcal{K}^{(2)}_{NM})\exp\left[-\frac{1}{2}\mathrm{tr}\{\hat{\Omega}^{-1}\hat{\theta}\tilde{\mathcal{K}}^{(2)}_{NN}\}\right]\\
&\equiv \exp\{\mathcal{L}^{(2)}_j\},\\
p(\tilde{y}_j|\beta_c, \delta_{cj}) &\geq \int \exp\{\mathcal{L}^{(1)}_{cj}\}p(u_j)p(b_j)du_j db_j\\
&= \mathcal{N}(\tilde{y}_j|\delta_{cj}X\beta_c, \hat{\Omega} + Z\hat{\Delta}Z^\top + \hat{\theta}\mathcal{K}^{(2)}_{NM}{K^{(2)}}^{-1}\mathcal{K}^{(2)}_{NM})\exp\left[-\frac{1}{2}\mathrm{tr}\{\hat{\Omega}^{-1}\hat{\theta}(\delta^2_{cj}\tilde{\mathcal{K}}^{(1)}_{NN} + \tilde{\mathcal{K}}^{(2)}_{NN})\}\right]\\
&\equiv \exp\{\mathcal{L}^{(2)}_{cj}\}.
\end{aligned}$$

The complete log likelihood of the mixture model is then written as

$$\mathcal{L} = \sum_{j=1}^{J} w_{0j}\mathcal{L}^{(2)}_j + w_{0j}\log \pi_0 + \sum_{c=1}^{C}\sum_{j=1}^{J}[w_{cj}\mathcal{L}^{(2)}_{cj} + w_{cj}\log \pi_c] + \sum_{c=1}^{C}\log p(\beta_c) + \sum_{c=1}^{C}\sum_{j=1}^{J}\log p(\delta_{cj}).$$

We use EM-algorithm to maximise the likelihood. The E-step computes

$$\begin{aligned}
\mathbb{E}_{\beta_c|\tilde{\mathcal{Y}}}\left[\mathcal{L}^{(2)}_{cj}\right] &= -\frac{N}{2}\log(2\pi) - \frac{1}{2}\log|V| - \frac{1}{2}(\tilde{y}_j - \delta_{cj}X\overline{\beta}_c)^\top V^{-1}(\tilde{y}_j - \delta_{cj}X\overline{\beta}_c)\\
&\quad - \frac{1}{2}\mathrm{tr}\left\{\hat{\Omega}^{-1}\theta(\delta^2_{cj}\tilde{\mathcal{K}}^{(1)}_{NN} + \tilde{\mathcal{K}}^{(2)}_{NN})\right\} - \frac{1}{2}\mathrm{tr}\left\{\delta^2_{cj}X^\top V^{-1}X\mathbb{V}_{\beta_c}\right\},\\
\mathbb{E}_{\delta_{cj}|\tilde{\mathcal{Y}}}\left[\mathcal{L}^{(2)}_{cj}\right] &= -\frac{N}{2}\log(2\pi) - \frac{1}{2}\log|V| - \frac{1}{2}(\tilde{y}_j - \overline{\delta}_{cj}X\beta_c)^\top V^{-1}(\tilde{y}_j - \overline{\delta}_{cj}X\beta_c)\\
&\quad - \frac{1}{2}\mathrm{tr}\left[\hat{\Omega}^{-1}\theta\left\{\mathbb{E}[\delta^2_{cj}|\tilde{\mathcal{Y}}]\tilde{\mathcal{K}}^{(1)}_{NN} + \tilde{\mathcal{K}}^{(2)}_{NN}\right\}\right] - \frac{1}{2}\mathrm{Var}(\delta_{cj}|\tilde{\mathcal{Y}})\beta_c^\top X^\top V^{-1}X\beta_c,
\end{aligned}$$

$$\begin{aligned}
\mathbb{E}_{\delta_{cj}|\tilde{\mathcal{Y}}}\left[\mathbb{E}_{\beta_c|\tilde{\mathcal{Y}}}\left[\mathcal{L}_{cj}^{(2)}\right]\right] &= -\frac{N}{2}\log(2\pi) - \frac{1}{2}\log|V| - \frac{1}{2}(\tilde{y}_j - \overline{\delta}_{cj}X\overline{\beta}_c)^\top V^{-1}(\tilde{y}_j - \overline{\delta}_{cj}X\overline{\beta}_c) - \frac{1}{2}\mathrm{Var}(\delta_{cj}|\tilde{\mathcal{Y}})\overline{\beta}_c^\top X^\top V^{-1}X\overline{\beta}_c \\
&\quad - \frac{1}{2}\mathrm{tr}\left[\hat{\Omega}^{-1}\theta\left\{\mathbb{E}[\delta_{cj}^2|\tilde{\mathcal{Y}}]\tilde{\mathcal{K}}_{NN}^{(1)} + \tilde{\mathcal{K}}_{NN}^{(2)}\right\}\right] - \frac{1}{2}\mathrm{tr}\left\{\mathbb{E}[\delta_{cj}^2|\tilde{\mathcal{Y}}]X^\top V^{-1}X\mathbb{V}_{\beta_c}\right\} \\
&\equiv \overline{\mathcal{L}}_{cj}^{(2)}, \\
\mathbb{E}[w_{cj}|\tilde{\mathcal{Y}}] &= \frac{\pi_c\exp\{\overline{\mathcal{L}}_{cj}^{(2)}\}}{\pi_0\exp\{\mathcal{L}_j^{(2)}\} + \sum_{c=1}^{C}\pi_c\exp\{\overline{\mathcal{L}}_{cj}^{(2)}\}} \\
&\equiv \overline{w}_{cj},
\end{aligned}$$

where $\mathbb{E}[\beta_c|\tilde{\mathcal{Y}}] = \overline{\beta}_c$, $\mathrm{Var}[\beta_c|\tilde{\mathcal{Y}}] = \mathbb{V}_{\beta_c}$, $\mathbb{E}[\delta_{cj}|\tilde{\mathcal{Y}}] = \overline{\delta}_{cj}$ and $\mathbb{E}[\delta_{cj}^2|\tilde{\mathcal{Y}}] = \mathrm{Var}(\delta_{cj}|\tilde{\mathcal{Y}}) + \overline{\delta}_{cj}^2$.

The M-step becomes

$$\begin{aligned}
\pi_c &= \frac{1}{J}\sum_{j=1}^{J}\overline{w}_{cj}, \\
\beta_c|\tilde{\mathcal{Y}} &\sim \mathcal{N}(\mathbb{V}_{\beta_c}X^\top V^{-1}\sum_{j=1}^{J}\overline{w}_{cj}\overline{\delta}_{cj}\tilde{y}_j, \mathbb{V}_{\beta_c}), \\
\delta_{cj}|\tilde{\mathcal{Y}} &\sim \mathcal{N}(\mathbb{V}_{\delta_{cj}}\overline{\beta}_c^\top XV^{-1}\tilde{y}_j, \mathbb{V}_{\delta_{cj}}),
\end{aligned}$$

where

$$\begin{aligned}
\mathbb{V}_{\beta_c} &= \left[\sum_{j=1}^{J}\overline{w}_{cj}\mathbb{E}[\delta_{cj}^2|\tilde{\mathcal{Y}}]X^\top V^{-1}X + (\theta K^{(1)})^{-1}\right]^{-1}, \\
\mathbb{V}_{\delta_{cj}} &= \frac{1}{\mathrm{tr}\left\{X^\top V^{-1}X(\mathbb{V}_{\beta_c} + \overline{\beta}_c\overline{\beta}_c^\top)\right\} + \mathrm{tr}\{\hat{\Omega}^{-1}\theta\tilde{\mathcal{K}}_{NN}^{(1)}\} + 1},
\end{aligned}$$

since

$$\begin{aligned}
\frac{\partial}{\partial\delta_{cj}}\mathbb{E}_{\beta_c|\tilde{\mathcal{Y}}}\left[\overline{w}_{cj}\mathcal{L}_{cj}^{(2)} + \log p(\delta_{cj})\right] &= \overline{\beta}_c^\top XV^{-1}\tilde{y}_j - \delta_{cj}\left[\mathrm{tr}\left\{X^\top V^{-1}X(\mathbb{V}_{\beta_c} + \overline{\beta}_c\overline{\beta}_c^\top)\right\} + \mathrm{tr}\{\hat{\Omega}^{-1}\theta\tilde{\mathcal{K}}_{NN}^{(1)}\} + 1\right], \\
\frac{\partial}{\partial\beta_c}\mathbb{E}_{\delta_{cj}|\tilde{\mathcal{Y}}}\left[\sum_{j=1}^{J}\overline{w}_{cj}\mathcal{L}_{cj}^{(2)} + \log p(\beta_c)\right] &= \sum_{j=1}^{J}\overline{w}_{cj}\overline{\delta}_{cj}X^\top V^{-1}\tilde{y}_j - \left[\sum_{j=1}^{J}\overline{w}_{cj}\mathbb{E}[\delta_{cj}^2|\tilde{\mathcal{Y}}]X^\top V^{-1}X + (\theta K^{(1)})^{-1}\right]\beta_c.
\end{aligned}$$

#### 1.5 GP regression for mapping eQTLs and Bayes factor calculation

The genetic association mapping model also uses the estimated model parameter of the GPLVM. Let $g_l$ denotes the vector of genotype dosages, the $i$th element of which indicates the number of alternative alleles of the genetic origin of cell $i$ at the biallelic genetic variant $l$. We model the genetic association as a gene-environment interaction between the genotype and the GP $f_{jl}$ governed by the kernel $\hat{\theta}\mathcal{K}_{NN}^{(1)}$ for the target cell state:

$$\begin{aligned}
y_j|f_{jl}, f_j, b_j &\sim \mathcal{N}(v_{jl}g_l + f_{jl}\odot g_l + W\hat{\mathcal{A}} + f_j + Zb_j, \sigma_{jl}^2\hat{\Omega}) \\
v_{jl} &\sim \mathcal{N}(0, \delta_g^2\sigma_{jl}^2) \\
f_{jl} &\sim \mathcal{N}(0, \delta_g^2\sigma_{jl}^2\hat{\theta}\mathcal{K}_{NN}^{(1)}) \\
f_j &\sim \mathcal{N}(0, \sigma_{jl}^2\hat{\theta}\mathcal{K}_{NN}) \\
b_j &\sim \mathcal{N}(\hat{\zeta}, \sigma_{jl}^2\hat{\Delta})
\end{aligned}$$

where $\delta_g^2$ is an arbitrary genetic variance parameter. For a rigorous association mapping, we the gene-specific residual $\sigma_{jl}$ as a free parameter.

In order to make the computation tractable, we assume

$$\begin{aligned} f_{jl}|u_{jl} &\sim \mathcal{N}(\mathcal{K}_{NM}{}^{(1)}K^{(1)}{}^{-1}u_{jl}, \delta_g^2\sigma_j^2\hat{\theta}\tilde{\mathcal{K}}_{NN}^{(1)}) \\ f_j|u_j &\sim \mathcal{N}(\mathcal{K}_{NM}K^{-1}u_j, \sigma_{jl}^2\hat{\theta}\tilde{\mathcal{K}}_{NN}) \\ u_{jl} &\sim \mathcal{N}(0, \delta_g^2\sigma_{jl}^2\hat{\theta}K^{(1)}) \\ u_j &\sim \mathcal{N}(0, \sigma_{jl}^2\hat{\theta}K) \end{aligned}$$

then a lower bound can be written as

$$\begin{aligned} \log p(y_j|v_{jl}, u_{jl}, u_j, b_j) &= \log \int p(y_j|f_{jl}, f_j, b_j)p(f_{jl}|\beta_c)p(f_j|u_j)df_{jl}df_j \\ &\geq \int \log p(y_j|f_{jl}, f_j, \delta_{jc}, b_j)p(f_{jl}|\beta_c)p(f_j|u_j)df_{jl}df_j \\ &= \log \mathcal{N}(y_j|v_{jl}g_l + \bar{f}_{jl} \odot g_l + W\hat{\mathcal{A}} + \bar{f}_j + Zb_j, \sigma_{jl}^2\hat{\Omega}) - \frac{1}{2}\mathrm{tr}\{\hat{\Omega}^{-1}\hat{\theta}(\delta_g^2 G_l\tilde{\mathcal{K}}_{NN}^{(1)}G_l + \tilde{\mathcal{K}}_{NN})\} \\ &\equiv \mathcal{L}_{jl}^{(1)}, \end{aligned}$$

where $G_l = \mathrm{diag}(g_l)$. We then arrive at the Titsias bounds:

$$\begin{aligned} p_0(y_j) &\geq \int \exp\{\mathcal{L}_{jl}^{(1)}|_{v_{jl}=0, u_{jl}=0}\}p(u_j)p(b_j)du_jdb_j \\ &= \mathcal{N}\left(y_j|W\hat{\mathcal{A}}, \sigma_j^2(\hat{\Omega} + Z\hat{\Delta}Z^\top + \hat{\theta}\mathcal{K}_{NM}K^{-1}\mathcal{K}_{NM})\right) \exp\left[-\frac{1}{2}\mathrm{tr}\{\hat{\Omega}^{-1}\hat{\theta}\tilde{\mathcal{K}}_{NN}\}\right] \\ &\equiv \exp\{\mathcal{L}_j^{(2)}\}, \\ p(y_j) &\geq \int \exp\{\mathcal{L}_{jl}^{(1)}\}p(v_{jl})p(u_{jl})p(u_j)p(b_j)du_{jl}du_jdb_j \\ &= \mathcal{N}\left(y_j|W\hat{\mathcal{A}}, \sigma_{jl}^2(\hat{\Omega} + Z\hat{\Delta}Z^\top + \hat{\theta}\mathcal{K}_{NM}K^{-1}\mathcal{K}_{NM} + \delta_g^2 g_l g_l^\top + \delta_g^2\hat{\theta}G_l\mathcal{K}_{NM}^{(1)}K^{(1)}{}^{-1}\mathcal{K}_{NM}^{(1)}G_l)\right) \\ &\quad\times \exp\left[-\frac{1}{2}\mathrm{tr}\{\hat{\Omega}^{-1}\hat{\theta}(\delta_g^2 G_l\tilde{\mathcal{K}}_{NN}^{(1)}G_l + \tilde{\mathcal{K}}_{NN})\}\right] \\ &\equiv \exp\{\mathcal{L}_{jl}^{(2)}\}. \end{aligned}$$

Therefore the Bayes factor of genetic association is obtained by

$$\begin{aligned} \log BF_{jl} &\equiv \mathcal{L}_{jl}^{(2)} - \mathcal{L}_j^{(2)} \\ &= -\frac{N}{2}\log(\hat{\sigma}_{jl}^2) + \frac{N}{2}\log(\hat{\sigma}_j^2) - \frac{1}{2}\log|\Delta_g^{-1} + Z_l^\top\hat{\Omega}^{-1}Z_l - Z_l^\top\hat{\Omega}^{-1}\tilde{Z}\Phi\tilde{Z}^\top\hat{\Omega}^{-1}Z_l| \\ &\quad - \frac{1}{2}\log|\Delta_g| - \frac{1}{2}\mathrm{tr}\{\hat{\Omega}^{-1}\hat{\theta}\delta_g^2 G_l\tilde{\mathcal{K}}_{NN}^{(1)}G_l\} \end{aligned}$$

where

$$\begin{aligned} \Delta_g &= \mathrm{diag}(\delta_g^2\theta K, \delta_g^2), \\ Z_l &= (G_l\mathcal{K}_{NM}{}^{(1)}K^{(1)}{}^{-1}, g_l), \\ V_l &= \hat{\Omega} + \tilde{Z}\tilde{\Delta}\tilde{Z}^\top + Z_l\Delta_g Z_l^\top, \\ \hat{\sigma}_{jl}^2 &= (y_j - W\hat{\mathcal{A}})_j^\top V_l^{-1}(y_j - W\hat{\mathcal{A}}), \end{aligned}$$

and

$$
\begin{aligned}
|V_l| &= |\text{diag}(\tilde{\Delta}, \Delta_g)^{-1} + (\tilde{Z}, Z_l)^\top \hat{\Omega}(\tilde{Z}, Z_l)||\hat{\Omega}||\text{diag}(\tilde{\Delta}, \Delta_g)| \\
&= |\Phi^{-1}||\Delta_g^{-1} + Z_l^\top \hat{\Omega}^{-1} Z_l - Z_l^\top \hat{\Omega}^{-1} \tilde{Z} \Phi \tilde{Z}^\top \hat{\Omega}^{-1} Z_l||\hat{\Omega}||\text{diag}(\tilde{\Delta}, \Delta_g)|.
\end{aligned}
$$
